# Large extracellular vesicles containing mitochondria (EVMs) derived from Alzheimer’s disease cells harbor pathologic functional and molecular profiles and spread mitochondrial dysfunctions

**DOI:** 10.1101/2025.01.08.631851

**Authors:** Fanny Eysert, Véronique Legros, Anne-Sophie Gay, Delphine Debayle, Sandra Lacas-Gervais, Khouloud Kaidi, Guillaume Chevreux, Julien Lagarde, Fréderic Checler, Marie Sarazin, Marie-Claude Potier, Mounia Chami

## Abstract

In addition to small extracellular vesicles known as exosomes, cells release large extracellular vesicles containing mitochondria (EVMs). However, the molecular and functional characteristics of EVMs, as well as the impact of EVM secretion on the spreading of mitochondrial dysfunction between cells, remain unknown in the context of Alzheimer’s Disease (AD). Here, we provide an ultrastructural, biochemical, and functional characterization of EVMs isolated from cells expressing the amyloid precursor protein (APP) with the familial Swedish mutation (APPswe). We identified differential proteomic and lipidomic signatures in APPswe-derived EVMs compared to control EVMs and revealed a specific proteomic profile in EVMs isolated from conditioned media of fibroblasts from AD patients at the prodromal stage of the disease. Our findings show that APP-C terminal fragments (APP-CTFs) pathogenic accumulation in cells potentiates EVM secretion through the budding of the plasma membrane. We lastly demonstrated that APP-CTFs loaded EVMs are active carriers of dysfunctional mitochondria that transfer mitochondrial pathology to naïve control recipient cells.

## INTRODUCTION

Alzheimer’s disease (AD) is a devastating neurodegenerative disease characterized by the progressive decline of cognitive functions. AD-related proteinopathy encompasses the accumulation of intracellular aggregates of Tau protein in the neurofibrillary tangles and of extracellular aggregates of a set of amyloid beta (Aβ) peptides in the senile plaques^1^. Aβ peptides are the byproducts of the cleavage of the amyloid precursor protein (APP) by the β-secretase generating a soluble sAPPβ fragment and a membrane-anchored β-C terminal fragment of 99 amino acids (aa) (β-CTF:C99). This latter is cleaved by the α-secretase in the non-amyloidogenic pathway to produce the α-CTF of 83 aa (α-CTF:C83), or by the γ-secretase in the amyloidogenic pathway to produce Aβ peptides^2^.

In parallel to toxic Aβ, our team and others uncovered the contribution of C99 fragment in AD pathophysiology, eliciting synaptic alterations and apathy-like behavior^3–5^. The accumulation of APP-CTFs affects endolysosomal and mitochondrial functions in an Aβ-independent manner and impacts general autophagy as well as the specific mitophagy pathways^5–10^.

Studies have unveiled an overlap between autophagy and endolysosomal dysfunctions and the release of small (30-150 nm) vesicles (EVs), commonly known as exosomes, into the extracellular space^11^. Small EVs secretion occurs mainly through the fusion of multivesicular bodies (MVBs), which contain intraluminal vesicles (ILVs), with the plasma membrane in an exocytic process^12^ and are now recognized to be mediators of cellular communication with critical roles in cell physiology and pathology including in AD^13^. It has been shown that small EVs containing programmed cell death 6-interacting protein (ALIX) and flotillin-1, are associated with Aβ plaques in post-mortem AD brains^14^, and play a crucial role in the dissemination of APP-derived proteins as well as pathological tau in AD^15, 16^. Accordingly, it has been shown that both Aβ and APP-CTFs are present in small EVs derived from AD post-mortem brains as well as from AD murine brains and AD neurons in culture^17–20^.

Beside small EVs, cells release large extracellular microvesicles (100 – 1000 nm) by budding and shedding directly from the plasma membrane^12^. A subset of these large extracellular vesicles contains mitochondria (EVMs) and are released by different cell types under physiological and pathological conditions^21, 22^. It has been proposed that the transfer of EVMs and of isolated mitochondria from cell-to-cell is a potential mode for mitochondrial content spreading in neurodegenerative diseases such as stroke and in Down syndrome^23–25^. However, the molecular and the functional characterization of EVMs have not been described in AD. Moreover, the impact of EVMs secretion and on the spreading of mitochondria dysfunctions between cells is still unknown in AD.

We designed the current study to investigate if aberrant accumulation of dysfunctional mitochondria in AD cells, driven by APP-CTFs accumulation, would trigger the release of EVMs acting as damage molecules and spreading mitochondrial pathology. We used optimized protocols for the isolation and the functional characterization of EVMs obtained from conditioned media of control neuroblastoma cells or stably expressing the APP familial Swedish mutation (APPswe). We studied the proteomic and the lipidomic signatures of AD-derived EVMs versus exosomes and unveiled a specific proteomic signature in EVMs isolated from the conditioned media of fibroblasts isolated from sporadic AD patients at the prodromal stage of the disease. We investigated the impact of APP-CTFs accumulation on EVMs secretion and mitochondrial function, demonstrating, first, that EVMs secretion occurs through plasma membrane budding in an APP-CTFs-dependent manner, and second, unlike exosomes, EVMs secretion is not linked to autophagosome-lysosome dysfunction. We also provided evidences that AD EVMs propagate mitochondria structure and function alterations to naïve control cells.

Overall, our study demonstrates that AD-derived EVMs are active cargo harboring specific proteomic and lipidomic signatures propagating mitochondrial dysfunction and likely contributing to AD pathogenesis.

## RESULTS

### EVMs isolation and study of their ultrastructure, size and molecular content

We first established the experimental design of the study (Fig. 1A). We adapted a unique protocol allowing the isolation, of small extracellular vesicles named here after “exosomes” and of large vesicles identified here after as “extracellular vesicles containing mitochondria: EVMS” (Fig. 1A, B). Exosomes and EVMs were obtained from conditioned media depleted of serum (CM) of neuroblastoma cells expressing empty pcDNA3.1 vector (control: CTRL) or the amyloid precursor protein with the familial AD (FAD) double Swedish mutation (APPswe: APP K670M-N671L). We first validate our protocol for the isolation of EVMs by transmission electron microscopy (TEM) showing a heterogeneous population of vesicles differing in electron density where mitochondria cristae ultrastructure was easily visualized (Fig 1C). TEM demonstrate that EVMs contain both isolated mitochondria and mitochondria surrounded by a double membrane with approximative sizes ranging from 100 to 500 nm (Fig 1C). Nanoparticles Tracking Analysis (NTA) unveiled as expected larger size distribution of EVMs (mean size ranging between 120 to 240 nm) as compared to exosomes (mean size ranging between 80 and 120 nm) (Fig. 1D, E). NTA also reveal that EVMs are secreted to a lesser extent than exosomes (almost 1 log difference) (Fig 1F), and that exosomes and EVMs size and concentration did not differ between CTRL and APPswe conditions (Fig. 1E, F). Flux cytometry combined with the specific mitochondrial MitoTracker Green probe (MitoT) staining demonstrate that 80-90% of CTRL or APPswe-derived EVMs are MitoT positive (Fig. 1G). As controls we showed that the fraction depleted in EVMs (ΔEVMs) show only 20% of MitoT positive particles likely corresponding to remaining EVMs in this fraction (Fig.1 H). We used unstained EVMs and ΔEVMs (Fig. 1I, J) to set-up the background and to refine the gating strategy. The quantification of MitoT positive particles does not reveal any change between control and APPswe EVMs (Fig. 1K) supporting our finding with NTA (Fig. 1F). Complementary controls were used to ensure the reliability of our gating (supplementary Fig. 1A-F). These include: i) filtered PBS in the presence or absence of MitoT (supplementary Fig. 1A, B), ii) control- and APPswe-derived mitochondrial fraction (MitoFr) stained or not with MitoT (supplementary Fig. 1C, D), iii) control- and APPswe-derived EVMs and MitoFr stained or not with MitoT and permeabilized with Triton X-100 (supplementary Fig. 1E, F). In an alternative protocol, we stained EVMs with both MitoT and Annexin V (Ann V), a high affinity ligand for the phospholipid, shown to cloak phosphatidylserine residues externalized on the surface of large EVs^12^. We validated the duplexing of MitoT and Ann V staining by analyzing EVMs simply stained with MitoT or Ann V (supplementary Fig. 1G-J). The double staining with MitoT and Ann V unveiled that 45 % of CTRL EVMs and 47 % of APPswe EVMs are MitoT^+^ and Ann V^-^ (supplementary Fig. 1K-O) likely corresponding to isolated mitochondria. We also identified 42 % of CTRL EVMs and 43 % of APPswe EVMs are MitoT^+^ and Ann V^+^ (supplementary Fig. 1K-O) likely representing mitochondria surrounded with plasma membrane. These results support our observations by TEM (Fig. 1C). The MitoT^-^ and Ann V^-^ population (less than 10%) may correspond to background particles also detected in EVMs stained with MitoT alone (Fig. 1G).

**Figure 1:**
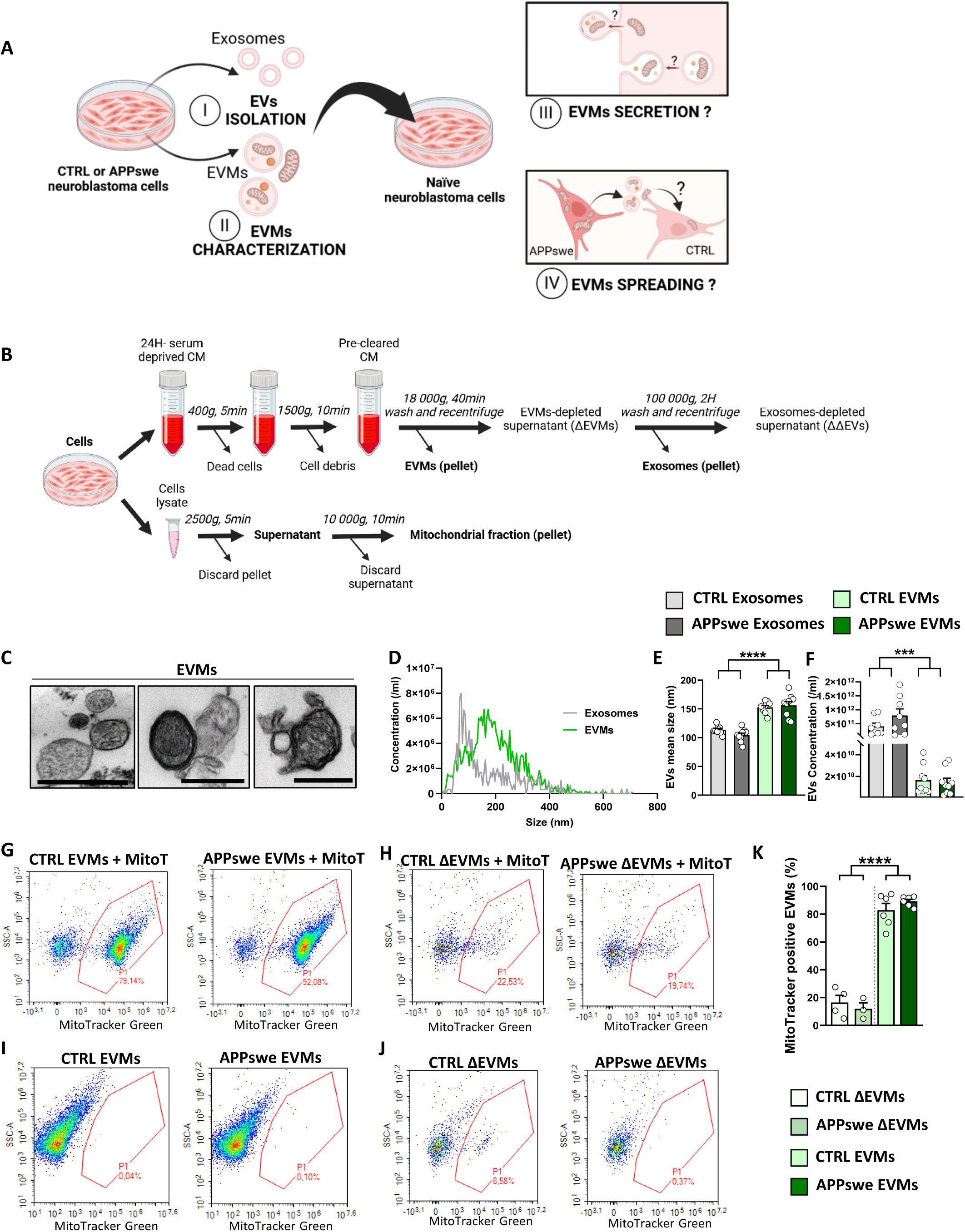
Study design, isolation and characterization of the large vesicles containing mitochondria (EVMs). **A)** The experimental design of the study includes: (I) the isolation and the comparative characterization of EVMs versus exosomes (EVs) obtained from serum-deprived conditioned media (CM) of control (CTRL) neuroblastoma cells and expressing the familial APPswe mutation (APPswe), (II) the comparative study of EVMs obtained from CTRL and APPswe cells, (III) the study of EVMs secretion, and (IV) the study of EVMs spreading. **B)** Schematic representation of the protocol used to isolate EVMs and exosomes. Cellular mitochondrial fraction was used as a control. **C)** Representative ultrastructures of EVMs obtained from CTRL cells using transmission electron microscopy. Scale bar: 200µm. **D)** Representative size distribution and concentration of control EVMs and exosomes (EVs) determined by Nanoparticles Tracking Analysis (NTA). **E-F)** Quantitative graphs of EVs mean size (E) and mean concentration/ml obtained from the same number of plated cells and quantified by NTA. **G, H)** Representative flow cytometry plots of EVMs (G) and ΔEVMs (H) stained with Mito tracker Green (MitoT). **I, J)** Representative flow cytometry plots of unstained EVMs (I) and ΔEVMs (J) used as controls for the gating strategy. **K)** Percentage of MitoT positive EVMs isolated from CTRL or APPswe cells. Graphs represent the mean ± SEM of n=9 individual preparations per condition (E, F), and n=6 or n= 4 individual preparations of EVMs or ΔEVMs (K), respectively. Statistical significance was determined using Two-way ANOVA (E, F) and Mann-Whitney test (K). ***, p<0,001; ****, p<0,0001. ns: not significant.

Altogether, these data validate the protocol we used for the isolation of EVMs. TEM, NTA and flux cytometry experiments concord to demonstrate that CTRL- and APPswe-derived EVMs similarly deliver isolated and plasma membrane-encapsulated mitochondria with adverse size larger than exosomes.

### Comparative biochemical and proteomic load of EVMs versus exosomes derived from control and APPswe cells

Then, we used SDS-PAGE to comparatively analyze biochemical signatures of EVMs and exosomes isolated form control and APPswe cells (Fig. 2A-I). We loaded exosomes and EVMs protein extracts obtained from the same number of cells. As expected, we show that control and APPswe exosomes express the canonical markers (Alix, CD63, CD81 and Flotillin-2) with a a non-significant reduction of Alix expression in APPswe exosomes versus CTRL ones (Fig. 2A-E). In addition, we reveal that control and APPswe EVMs are depleted of exosomal markers excepting Flotillin-2 and that they are enriched with mitochondria markers (TOMM20, TIMM23, HSP60, and HSP10) (Fig. 2A, F-I). We also do not report significant change in the selected mitochondrial proteins expression between CTRL and APPswe-derived EVMs (Fig. 2A, F-I). To analyze in depth the proteomic profile of exosomes and EVMs, we performed a label-free Nano-HPLC-HRMS analysis. A total of 1861 and 2088 proteins with 3 or more unique peptides were respectively identified in exosomes and EVMs. We selected proteins present in at least 3 samples out of 4 in each condition (bold numbers) (Fig. 2J-L) yielding to identify 1168 unique proteins in exosomes and 1292 unique proteins in EVMs (Fig. 2J-L). Among the selected proteins, 862 were common to both EVs subtype, 306 were only detected in exosomes and 430 in EVMs (Fig. 2L). The Principal Component Analysis (PCA) distinctly identify two clusters corresponding to exosomes and EVMs (Fig. 2M). Unique proteins identified in exosomes or EVMs and common proteins in both EVs were processed on the Database for Annotation, Visualization and Integrated Discovery (DAVID) according to Kyoto Encyclopedia of Genes and Genomes (KEGG) pathways (Fig. 2N) and Gene Ontology analysis (Fig. 2O-Q). KEGG pathway analysis unravels that exosomes unique proteins are grouped into pathways relative to “ECM-receptor interaction”, “Focal adhesion” or “ribosome” and that EVMs unique proteins are grouped into neurodegenerative diseases (Amyotrophic lateral sclerosis, Parkinson’s disease, Huntington’s disease, Prion disease, and Alzheimer’s disease) (Fig. 2N). Furthermore, mitochondrial relative pathways such as “reactive oxygen species” or “oxidative phosphorylation” were enriched in EVMs (Fig. 2N). Common proteins identified in Exosomes and EVMs share pathways related to “neurodegenerative diseases”, “proteasome” and “ribosome” (Fig. 2N). The analyses of biological process (BP), cellular component (CC) and molecular function (MF) reveal an enrichment with mitochondrial relative processes in EVMs proteome as compared to exosomes proteome shown to be enriched with “extracellular exosomes signaling”, “RNA processing” or “cell adhesion” (Fig 2O-Q). Here also, exosomes and EVMs proteome also share common BP, CC and MF components such as “extracellular exosomes” or “RNA binding” (Fig 2O-Q).

**Figure 2:**
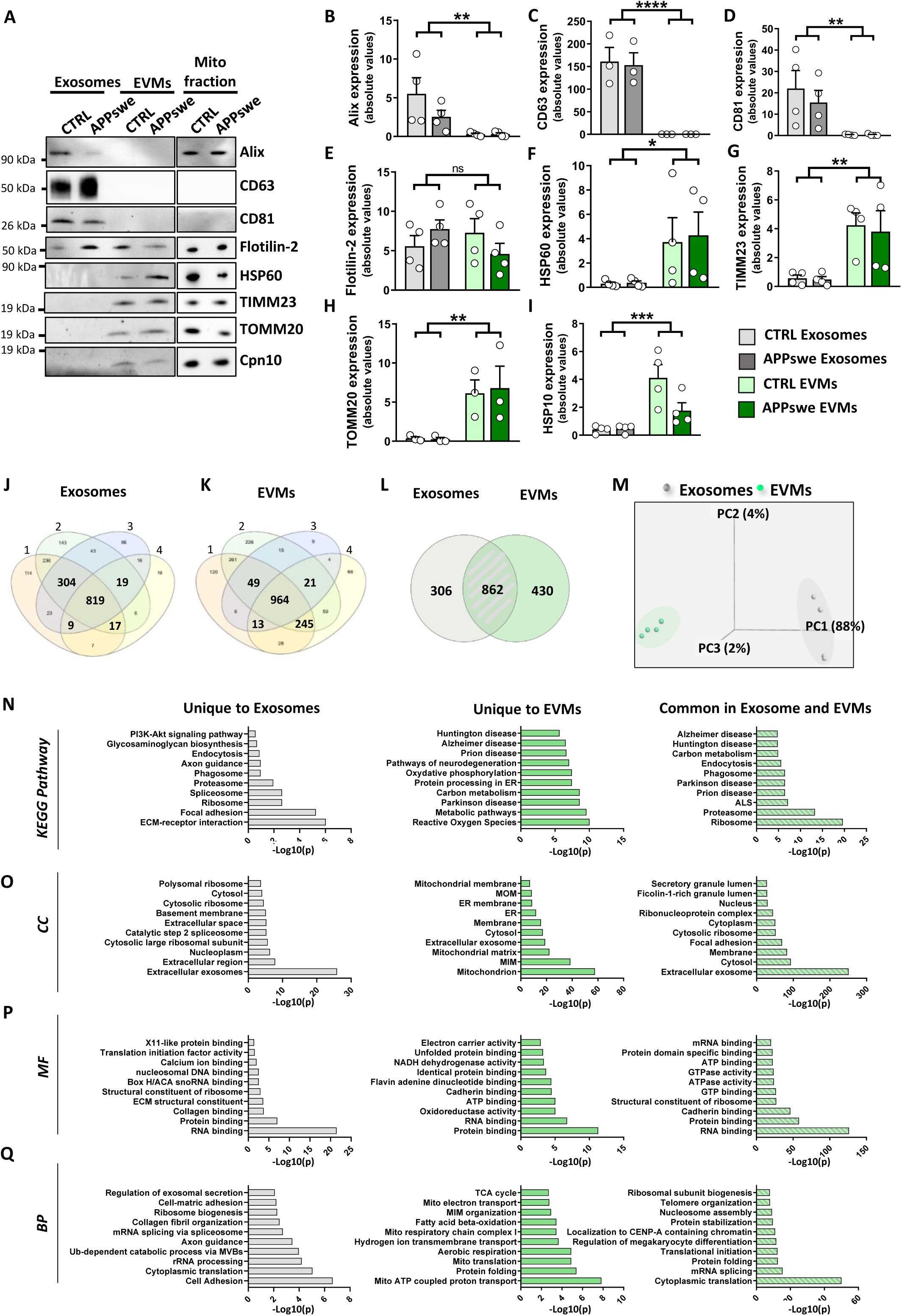
Biochemical and proteomic analyses demonstrating the enrichment of EVMs with mitochondrial proteins. **A-I)** SDS-page (A) and quantitative graphs of indicated proteins in exosomes and EVMs isolated from CM of CTRL or APPswe cells. The cellular mitochondrial fraction (Mito fraction) was loaded as a control for mitochondrial markers. **J-Q)** Comparative proteomic data of EVMs and exosomes. **J-K)** Venn diagram representing the number of proteins identified in each of the four replicates of exosomes (J) or EVMs (K) and detected with 3 or more unique peptides. **L)** Venn diagram representing the number of proteins identified in at least 3 out of 4 replicates of exosomes and EVMs. **M)** Principal component analysis (PCA) of four individual exosomes and EVMs preparations. **N)** KEGG Pathway, and **O-Q)** Gene Ontology (GO) analysis using the DAVID knowledgebase (v2023q2). The GO terms of top 10 Cellular Component (CC) (O), Molecular Function (MF) (P) and Biological Process (BP) (Q) of proteins identified in exosomes, EVMs or common in exosomes and EVMs. Abbreviations: ECM: extracellular matrix; Ub: ubiquitin; ER; endoplasmic reticulum; MOM: mitochondrial outer membrane; MIM: mitochondrial inner membrane; TCA: tricarboxylic acid; Mito: mitochondria; ALS: amyotrophic lateral sclerosis; Val: valine; Leu: leucine; Ile: isoleucine. (B-I) Graphs represent the mean ± SEM of n=3 or 4 individual preparations of EVMs and exosomes. (B-I) Statistical significance was determined by Two-way ANOVA *p <0,05, **p <0,01, ***p< 0,001, ****p<0,0001. The effect size between exosomes and EVMs measured by Hedges test is 1.52 (B), 3.356 (C), 1.581 (D), 0.467 (E), 1.04 (F), 2.555 (G), 2.178 (H) and 2.358 (I). (N-Q) Statistical significances correspond log-transformed p-values -log_10_ (p).

These biochemical and proteomic data converge to demonstrate that EVMs are largely enriched in specific mitochondrial signaling molecules.

### APPswe-derived EVMs accumulate APP-CTFs and harbor a differential proteomic signature versus CTRL-derived EVMs

Since previous studies have shown that secreted exosomes contain APP and the APP-CTFs^17–20^ and that we have showed that mitochondrial alterations in APPswe cells are triggered by APP-CTFs accumulation in mitochondria^10^ (Fig. 3A), we reasoned that APPswe-derived EVMs may similarly contain APP and APP-derived fragments. SDS-page analysis of EVs isolated from conditioned media of CTRL or APPswe cells, showed that both exosomes and EVMs derived from APPswe cells are positive for full-length APP and APP-CTFs (C83 and C99). However, we noticed that exosomes are largely enriched in APP-CTFs as compared to EVMs (Fig. 3A-D). Then, we analyzed in EVMs the sequence coverage of the identified APP N-terminal (N-ter) and APP C-terminal (C-ter) peptides respective to APP cleavage by α-, β-, δ-, and η-secretases, generating α-CTF (C83), β-CTF (C99), δ374-CTF, δ586-CTF, and η-CTF (Fig. 3E). This analysis reveals a large increase of both APP N-ter APP C-ter in APPswe-derived EVMs in comparison of control EVMs (Fig. 3F, G).

**Figure 3:**
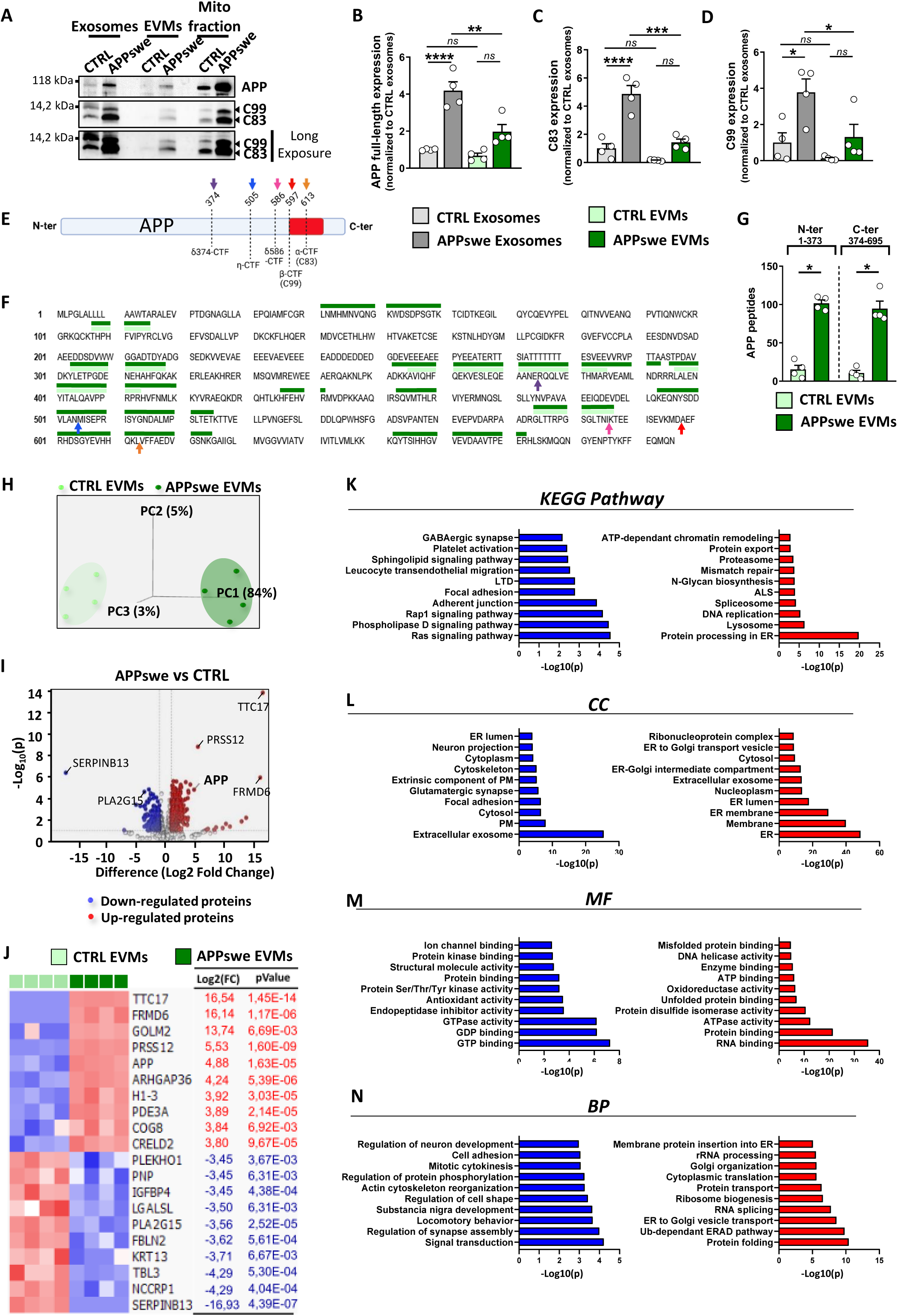
APPswe-derived EVMs accumulate APP and APP-CTFs and harbor a differential proteomic expression profile versus control EVMs. **A-D)** SDS page (A) and quantification of full-length APP (APP) (B) and APP-CTFs (C83 and C99) (C, D) in exosomes and EVMs isolated from CM of CTRL or APPswe cells. The cellular mitochondrial fraction (Mito fraction) was loaded as a control. **E)** Scheme of the full-length APP showing the cleavage sites (colored arrows) generating APP-αCTF, APP-βCTF, APP-ηCTF and two APP-δCTFs. **F)** Sequence coverage of APP-derived peptides identified by LC-MS/MS in EVMs obtained from CM of CTRL or APPswe cells (light green and dark green lines respectively). **G)** Quantification of the number of APP C-terminal (APP-Cter) and APP N terminal (APP-Nter) peptides obtained in CTRL or APPswe EVMs. **H)** Principal component analysis (PCA) of CTRL or APPswe EVMs obtained from n=4 individual preparations. **I)** Volcano plot of the log-transformed fold change versus log-transformed p-values (-log_10_(p)) of detected proteins. Gray dot lines indicate the cutoff of -Log_10_(p) >1 and Log2 fold change at -1 or +1 values. Blue dots represent the down-regulated proteins and red dots represent the up-regulated proteins in APPswe EVMs versus CTRL EVMs. **J)** Heatmap representation of the significant top 10 down-regulated (blue) and up-regulated (red) proteins in APPswe EVMs versus CTRL EVMs with filtering conditions of number of unique peptides >3, Log2(FC) <-1 or >+1 and p value <0,01. **K)** KEGG Pathway, and **L-N)** Gene Ontology (GO) terms of top 10 Cellular Component (CC) (L), Molecular Function (MF) (M) and Biological Process (BP) (N). Abbreviations: LTD: long-term depression; ER: endoplasmic reticulum; PM: plasma membrane; Ser: serine; Thr: threonine; Tyr: tyrosine; ALS: amyotrophic lateral sclerosis; Ub: ubiquitin. Graphs represent protein mean expression levels ± SEM of at least 4 individual preparations of exosomes and EVMs. B-D) Statistical significance was determined by One-way ANOVA test *p<0,05; **p<0,01 (B-D). The effect size measured by the Hedges test between CTRL and APPswe exosomes is 4.101 (B), 3.481 (C) and 1.193 (D). The effect size between CTRL and APPswe EVMs is 1.914 (B), 3.181 (C) and 1.037 (D). (K-N) Statistical significances correspond to log-transformed p-values (-log_10_ (p).

We further analyzed the proteome of individual preparations of CTRL- and APPswe-derived EVMs, identifying a total of 2088 proteins in CTRL-EVMS and 3238 proteins in APPswe-EVMs with 3 or more unique peptides. Among them, we selected proteins present in at least 3 samples out of 4 in each condition (bold numbers) (supplementary Fig. 2A, B). Although we used similar proteins amount to perform proteomic analyses, we intriguingly identified a larger proteome content in APPswe-EVMs (2172 proteins) versus CTRL-EVMs (1292 proteins). Among them, 1212 proteins were commonly detected in control-EVMs and APPswe-EVMs, 80 proteins were detected only in CTRL-EVMs, and 960 proteins were detected only in APPswe-EVMs (supplementary Fig. 2B). We accordingly processed unique and common proteins using DAVID KEGG pathways (supplementary Fig. 2C) and Gene Ontology analysis (supplementary Fig. 2D-F). The most significant difference was an enrichment in “metabolism” pathways and in organelles “Endoplasmic Reticulum, Mitochondria and Nucleus” CC in APPswe-EVMs versus CTRL-EVMs (supplementary Fig. 2D-F). The quantitative PCA showed two distinct clusters representing CTRL-EVMs and APPswe-EVMs (Fig. 3H). We found a substantial number of significantly down- or up-regulated proteins, including APP in the top 10 of upregulated proteins in APPswe-EVMs versus CTRL-EVMs (Fig. 3I, J). The analysis of KEGG pathways (Fig. 3K) and of Gene Ontology (Fig. 3L-N) unraveled specific proteomic signatures of down-regulated (blue) or up-regulated (red) proteins with a significant upregulation in APPswe-EVMs of proteins linked to organelles CC and MF such as “Endoplasmic Reticulum, Mitochondria and Golgi-apparatus” (Fig. 3L-M).

### APPswe-derived EVMs harbor altered mitochondria proteomic signature

As APP-CTFs accumulation is known to disrupt mitochondria function, we underwent an in-depth proteomic analysis of mitochondrial proteins in EVMs by crossing the list of significant up- and down-regulated total proteins with the MitoCarta3.0 protein list^26^. We used the list of the identified mitochondrial-related proteins in EVMs for further analysis. The PCA first showed two distinct clusters representing CTRL EVMs and APPswe EVMs (Fig. 4A), demonstrating the differential mitochondrial proteins signature between both conditions. Quantitative analysis highlighted down- and up-regulated mitochondrial proteins in APPswe EVMs versus CTRL EVMs as shown in the volcano plot (Fig. 4B). The obtained heatmap revealed significant (p<0.01) down- (6) and up-regulated (39) proteins in APPswe EVMs versus CTRL EVMs (Fig. 4C). We then mapped the protein-protein interaction network of significant up- or down-regulated proteins according to their biological process. This molecular profiling highlighted mitochondrial alterations in mitochondrion organization, reactive oxygen species (ROS) metabolic process, aerobic respiration, lipid and nitrogen metabolic processes as well as mitochondrial RNA metabolic process (Fig. 4D). The KEGG pathway (Fig. 4E) and GO analysis (Fig. 4F, G) highlight several functions of mitochondria differentially expressed in APPswe-EVMs versus CTRL-EVMs consolidating the altered pathways identified by STRING analysis.

**Figure 4:**
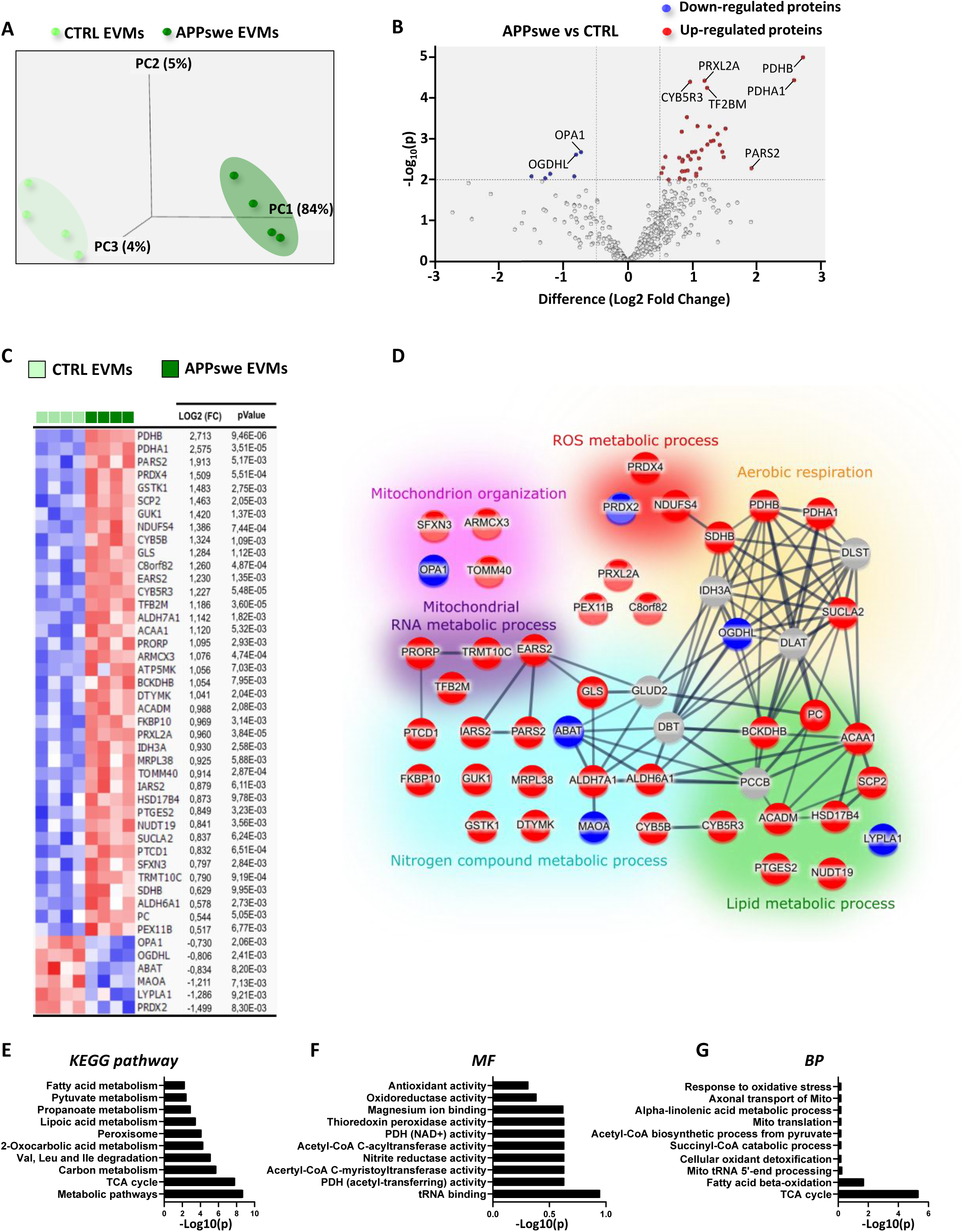
APPswe-derived EVMs show a differential mitochondrial proteomic signature versus control EVMs. **A)** Principal component analysis (PCA) of mitochondrial proteins from n=4 individual APPswe or CTRL EVMs preparations. **B)** Volcano plot of the log-transformed fold change versus log-transformed p-values (-log_10_ (p)) of detected proteins. Gray dot lines indicate the cutoff of -Log_10_(p) >2 and Log2 fold change at -0.5 or +0.5 values. Blue dots represent the down-regulated proteins and red dots represent the up-regulated proteins in APPswe EVMs versus CTRL EVMs. **C)** Heatmap representation of the significant down-regulated (blue) and up-regulated (red) proteins in APPswe EVMs versus CTRL EVMs with filtering conditions of number of unique peptides >3, Log2(FC) <-0,5 or >+0,5 and p value <0,01. **D)** Functional protein-protein interactome and high-confidence network analysis (STRING) (interaction score > 0.7) illustrating down-regulated (dark blue) and up-regulated (red) proteins in APPswe EVMs versus CTRL EVMs. Non-regulated proteins are in grey. **E)** KEGG pathway, and **F,G)** Gene Ontology (GO) terms of top 10 Molecular Function (MF) (F) and Biological Process (BP) (G) with -log_10_ (pValue) of significance mitochondrial proteins differentially expressed in APPswe versus CTRL-derived EVMs. Abbreviations: Val: valine; Leu: leucine; Ile: isoleucine; TCA: tricarboxylic acid; PDH: pyruvate dehydrogenase; Mito: mitochondria.

### EVMs isolated from fibroblasts of AD patients with mild-cognitive impairment present altered mitochondrial proteomic signature

Extracellular vesicles (EVs) have garnered significant attention as potential biomarkers for AD due to their ability to carry molecular cargo that can serve as disease biomarkers^27^. Interestingly, we and others have recently demonstrated that fibroblasts from AD patients harbor mitochondrial structure and function alterations as well as mitophagy failure phenotype^28–30^, mimicking the mitochondrial dysfunctions observed in AD neurons^8–10^. Since EVMs are secreted by different cell types including fibroblasts, we thought to comparatively study proteomic cargo of EVMs isolated from CM of primary fibroblasts isolated from previously studied control individuals (CTRL) (n=4) and sporadic AD (SAD) patients exhibiting either mild-cognitive impairments (AD-MCI) (n=6) or severe dementia (AD-D) (n=4)^29^ (Supplementary table 1 and Fig. 5A). We followed the experimental protocol for EVMs preparation described above (Fig. 1B) and restrain our study the analyzes of protein content of EVMs. We identified a total of 1842 proteins in CTRL EVMs, 1947 proteins in AD-MCI EVMs and 1655 proteins in AD-D EVMs with 3 or more unique peptides (supplementary Fig. 3 A-C). We selected proteins present in at least 3 samples out of 4 in CTRL and AD-D fibroblasts-derived EVMs (bold numbers) or proteins present in at least 5 out of 6 in AD-MCI fibroblast-derived EVMs (supplementary Fig. 3A-C). This filtering leads us to select a total of 1030 proteins in CTRL EVMs, 1125 proteins in AD-MCI EVMs and 1011 proteins in AD-D EVMs. We identified 879 common proteins in the three groups of EVMs, 71 unique proteins in CTRL EVMs, 118 unique proteins in AD-MCI EVMs, and 38 unique proteins detected in AD-D EVMs (Fig. 5B). Here also the PCA indicated three distinct clusters representing CTRL, AD-MCI and AD-D EVMs (Fig. 5C). Unique as well as the common proteins in the three groups of EVMs were processed on DAVID according to KEGG pathways (Fig. 5D and supplementary Fig. 3D) and GO analysis (Fig. 5E-G and supplementary Fig. 3E-G). Interestingly, we noticed an enrichment of mitochondrial relative pathways as well as an enrichment of mitochondrial components, molecular function and biological process in AD-MCI EVMs as compared to CTRL or AD-D EVMS (Fig. 5D-G). We then quantitatively assessed the proteins significantly up- or down-regulated in AD-D versus CTRL, in AD-MCI versus CTRL and in AD-D versus AD-MCI fibroblasts-derived EVMs (supplementary Fig. 3H-J) and filtered the proteins list according to the MitoCarta3.0 protein list^26^. This analysis showed 28 significant up- and 2 down-regulated proteins in AD-MCI versus CTRL fibroblasts-derived EVMs (Fig. 5H, I) and 24 significant up- and 1 down-regulated in AD-MCI versus AD-D fibroblasts-derived EVMs (Fig. 5J, K) while only one protein was up-regulated in AD-D versus CTRL fibroblasts-derived EVMs (Fig. 5L). Mitochondrial proteins of AD-MCI vs CTRL EVMs and of AD-D vs AD-MCI EVMs were then mapped into a STRING network according to their biological process. Intriguingly, we highlighted a specific upregulation of molecular components implicated in mitochondria organization, nitrogen compound metabolic process and aerobic respiration in AD-MCI EVMs but not in AD-D EVMs (Fig. 5I, K). EVMs mitochondrial proteome signature mimics the pathways identified in APPswe EVMs suggesting common mitochondrial proteomic signature in FAD-like EVMs and SAD-derived EVMs.

**Figure 5:**
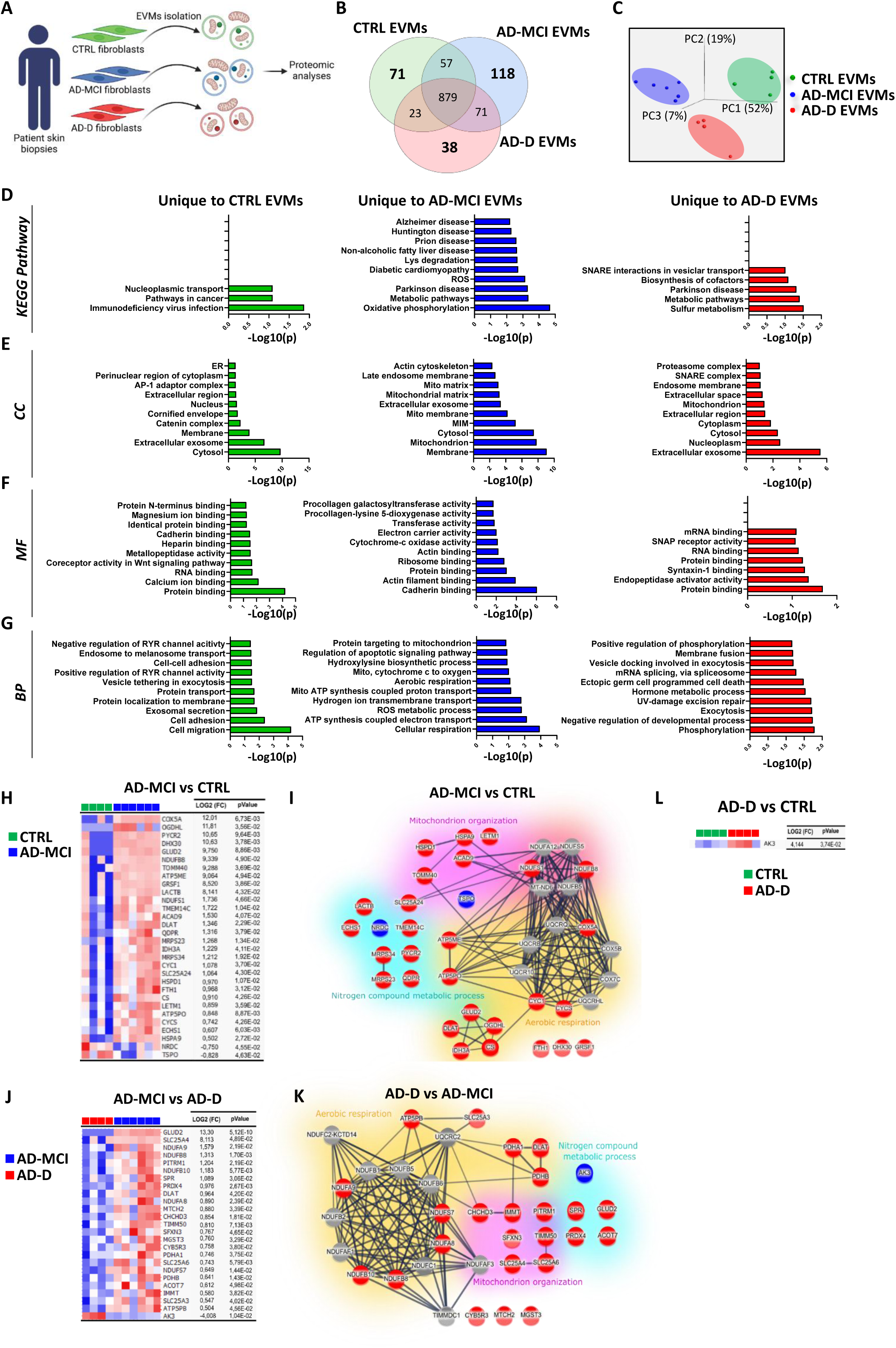
AD-MCI fibroblasts-derived EVMs harbor a specific mitochondrial signature. **A)** The experimental design includes the isolation and the comparative characterization of EVMs obtained from conditioned media of human fibroblasts healthy individuals (CTRL) or AD patients with mild-cognitive impairment (AD-MCI) or severe dementia (AD-D). **B)** Venn diagram representing the number of proteins identified in EVMs obtained from CTRL, AD-MCI or AD-D fibroblasts. **C)** Principal component analysis (PCA) of EVMs obtained from CTRL fibroblasts (n=4), AD-MCI fibroblasts (n=6) and AD-D fibroblasts (n=4). **D)** KEGG Pathway, and **E-G)** Gene Ontology (GO) terms of top 10 Cellular Component (CC) (E), Molecular Function (MF) (F) and Biological Process (BP) (G). Abbreviations: ER: endoplasmic reticulum; RYR: ryanodine receptor; Lys: lysine; ROS: reactive oxygen species; Mito: mitochondria; MIM; mitochondrial inner membrane; UV: ultra-violet. **H, J, L)** Heatmaps representation of the significant down-regulated (blue) and up-regulated (red) mitochondrial proteins in EVMs isolated from conditioned media (CM) of AD-MCI versus CTRL fibroblasts (H), AD-MCI versus AD-D fibroblasts (J), or AD-D versus CTRL fibroblasts (L). We applied filtering conditions of number of unique peptides >1, Log2(FC) <-0,5 or >+0,5 and p value <0,05. **L, K)** Functional protein-protein interactome and high-confidence network analysis (STRING) (interaction score > 0,7) illustrating down-regulated (dark blue) and up-regulated (red) proteins in AD-MCI EVMs versus CTRL EVMs (I) or AD-D EVMs versus AD-MCI EVMs. Non-regulated proteins are in grey. (D-G) Statistical significances correspond to log-transformed p-values (-log_10_ (p).

### APP-CTFs accumulation trigger an alteration of mitochondria function of EVMs

In addition to the identification of mitochondrial pathways differentially regulated in APPswe EVMs versus CTRL EVMs, we studied the impact of the accumulation of APP-CTFs, independently from Aβ peptides, on the function of mitochondria contained in EVMs. To this aim, we pharmacologically inhibited γ-secretase, thus blocking Aβ production and enhancing APP-CTFs levels as we previously reported^10^. We quantified the level of full-length APP and of APP-CTFs (C83 and C99) in both EVMs and used the cellular mitochondrial fractions of CTRL and APPswe cells as controls (Fig. 6A-D). The obtained results indicated an increase of APP-CTFs levels that reaches significance for C83 and C99 fragments between CTRL EVMs and APPswe EVMs isolated from the aforementioned cells treated with the γ-secretase inhibitor and for C99 fragment between EVMs isolated from APPswe cells treated with the γ-secretase inhibitor versus EVMs isolated from APPswe untreated cells (Fig. 6A-D). We analyzed mitochondrial membrane potential (ΔΨmit) in EVMs using TMRM probe. We observed a significant decrease of mitochondrial membrane potential in EVMs isolated from APPswe cells as compared to CTRL cells as well as in EVMs isolated from CTRL or APPswe cells treated with the γ-secretase inhibitor versus untreated cells (Fig. 6E). Moreover, the quantification of mitochondrial ROS (mitROS) levels revealed a significant increase within EVMs accumulating APP-CTFs (CTRL and APPswe) as compared to EVMs isolated from untreated cells (Fig. 6F). Altogether, these results demonstrate that EVMs accumulating APP-CTFs show mitochondrial membrane depolarization and increased ROS content.

**Figure 6:**
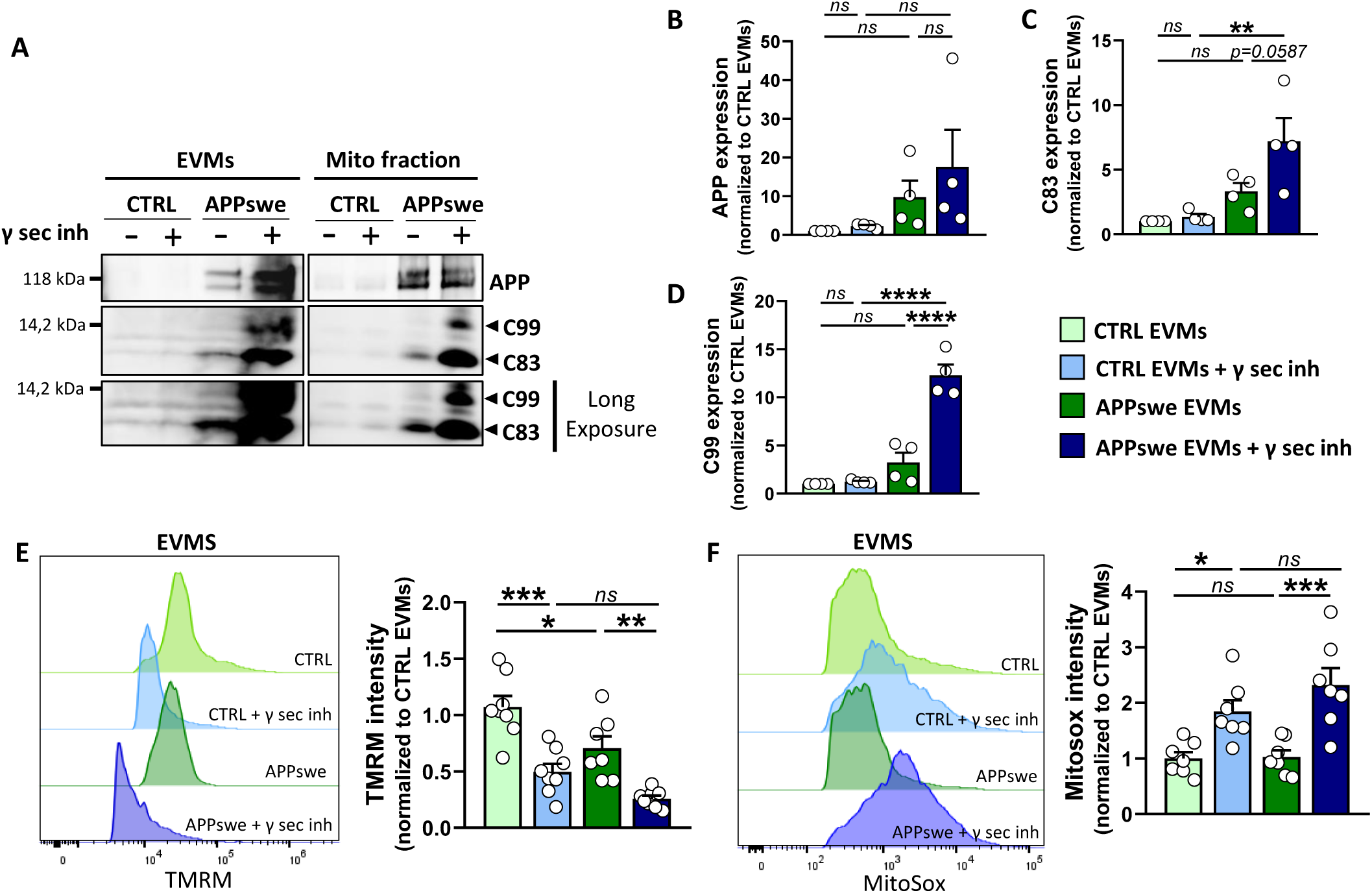
APPswe-derived EVMs show dysfunctional mitochondria. **A-D)** SDS page (A) and quantification (B-D) of full-length APP (B) and APP-CTFs (C83 and C99) (C,D) in EVMs isolated from CTRL or APPswe cells treated or not with the γ-secretase inhibitor (γ-sec inh). **E, F)** Representative flux cytometry profiles and quantitative graphs of fluorescence mean intensity of TMRM (E) and MitoSOX (F) of EVMs isolated from CTRL or APPswe cells treated or not with the γ-secretase inhibitor (γ sec inh). Graphs represent mean ± SEM of the expression levels of the indicated proteins (B-D) or of the fluorescence intensities of TMRM or MitoSOX obtained from n=4 (A-D), n=8 (E) or n=7 (F) independent preparations per condition. (B-F) Statistical significance was determined by two-way ANOVA test. *p<0,05; **p<0,01; ***p<0,001; ****p<0,0001. B-D) The effect size measured by the Hedges test between CTRL and APPswe exosomes is 2.585 (B), 0.926 (C) and 1.731 (D). The effect size between CTRL and APPswe EVMs is 0.461 (B), 1.247 (C) and 3.707 (D).

### APP-CTFs accumulation increases EVMs genesis and secretion through the budding of the plasma membrane

Recent studies have reported that mitochondria-derived vesicles (MDVs) are targeted to the late endosomes or MVBs in order to be degraded by the lysosomes or are redirected towards the plasma membrane for exocytosis^21, 31^. In this setting, we assessed in CTRL and APPswe cells treated or not with the γ-secretase inhibitor, the colocalization of mitochondria with CD63 positive compartments corresponding to MVBs as well as multivesicular endosomes, amphisomes or autophagosomes involved in exocytosis. To investigate the relationship between autophagic-lysosomal impairment and EVMs secretion, cells were treated with the vacuolar H+ V-ATPase, bafilomycin A1 (Baf A1) (Fig. 7A). In accordance with lysosomal defect previously reported in APPswe cells and shown to be linked to APP-CTFs accumulation^5^, we reported a significant increase of the number of CD63 positive puncta in APPswe cells as compared to CTRL cells (Fig. 7B). The number of CD63 positive puncta was also higher in γ-secretase and Baf A1-treated CTRL cells and between CTRL and APPswe cells treated with Baf A1 (Fig. 7B). In parallel, we noticed enhanced colocalization of APP-CTFs with CD63 in Baf A1 treated CTRL and APPswe cells and in APPswe cells versus CTRL cells treated with γ-secretase inhibitor (supplementary Fig. 4A, B). We accordingly report an enhanced size of CD63 puncta in CTRL cells treated with Baf A1 and in APPswe cells treated with either γ-secretase inhibitor or Baf A1, attesting to MVB enlargement and defective degradation. However, we observed a significant decrease in the colocalization between CD63+ organelles and mitochondria in untreated APPswe cells compared to CTRL cells, as well as following treatment with the γ-secretase inhibitor (Fig. 7C). These results demonstrate that, despite the increased number of CD63+ compartments associated with APP-CTF accumulation, mitochondria are not engulfed in CD63+ compartments for subsequent degradation or exocytosis. As described for microvesicles formation, MDVs could bud directly from the plasma membrane into extracellular space to generate EVMs^32^. The 3D reconstruction of confocal images (4 z-stacks focal planes) of the mitochondria stained with mitoTracker green and the plasma membrane stained with Wheat Germ Albumin showed a larger colocalization of mitochondria organelles with the plasma membrane in APPswe cells treated with the γ-sec inhibitor (indicated by black arrows in representative plots) as compared to CTRL cells treated with the γ-sec inhibitor (Fig. 7D). The proximity of mitochondria with the plasma membrane was almost undetectable in untreated conditions and in CTRL and APPswe cells treated with Baf A1 (Fig. 7D). The in-depth analyses of the 3D-reconstructed images revealed the presence of intracellular vesicles containing mitochondria (Fig. 7E, F) and plasma membrane blebs containing mitochondria (Fig. 7E, G). The quantification of these structures showed a significant increase in plasma membrane blebs containing mitochondria in APPswe cells treated with the γ-secretase inhibitor compared to both CTRL cells, treated or untreated with the γ-secretase inhibitor (Fig. 7E, G). However, the addition of Baf A1 significantly increased the number of intracellular vesicles containing mitochondria in both CTRL and APPswe cells (Fig. 7E, F). Accordingly, NTA unveiled a significant increase of EVMs secretion from APPswe cells treated with the γ-sec inhibitor, but not upon Baf A1 treatment (Fig. 7H).

**Figure 7:**
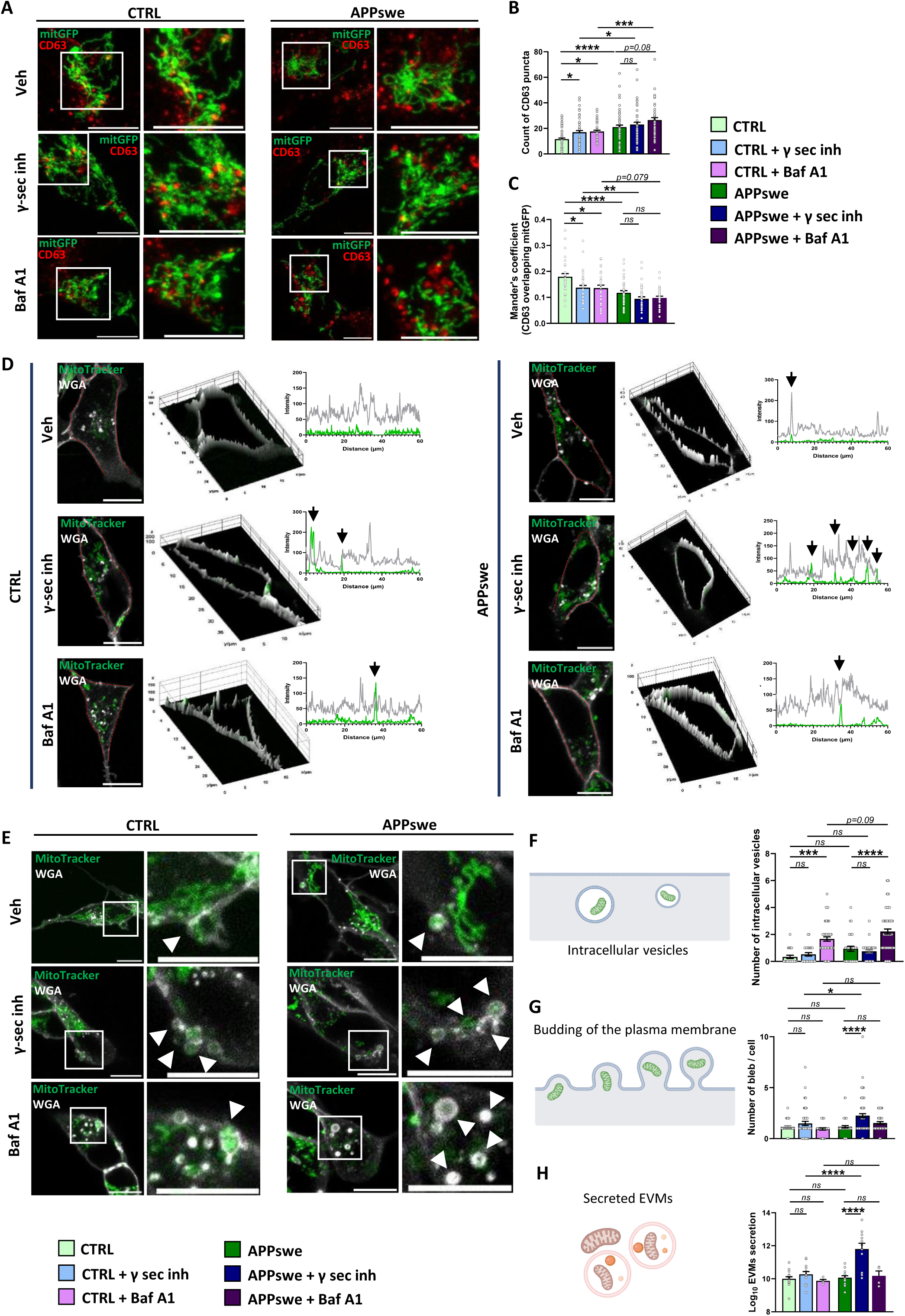
APP-CTFs accumulation increases the genesis and secretion of EVMs. **A)** Representative images of CTRL and APPswe cells treated with either vehicle (veh) used as control, the γ-secretase inhibitor (γ-sec inh) or bafilomycin A1 (Baf A1). Cells were transiently transfected with the mitochondrial probe mit-GFP (green) and stained with CD63 (red). The merged signal is shown in yellow. The high-magnification images correspond to the boxed areas. Scale bars = 10μm. **B)** Quantification of CD63-positive puncta from CTRL (n=80 cells), CTRL+γ-sec inh (n=66 cells), CTRL+Baf A1 (n=49 cells), APPswe (n=60 cells), APPswe+γ-sec inh (n=52 cells) and APPswe+Baf A1 (n=50 cells). **C)** Mander’s coefficient of mitGFP colocalization with CD63 from CTRL (n=34 cells), CTRL+γ-sec inh (n=39 cells), CTRL+Baf A1 (n=28 cells), APPswe (n=36 cells), APPswe+γ-sec inh (n=37 cells) and APPswe+Baf A1 (n=31 cells). **D)** Representative images of CTRL and APPswe cells treated as in (A) and co-stained with the membrane marker Wheat Germ albumin (WGA) (white) and the mitochondrial probe MitoTracker (green). The corresponding 3D representation of cells and profile plot using line scan analysis (dotted red line) show the colocalization of mitochondria within the plasma membrane (arrows). **E)** Representative images of CTRL and APPswe cells treated and stained as in (D). The high-magnification images correspond to the boxed areas. **F, G)** Schemes and quantifications of intracellular mitochondrial “vesicles” (F) and of plasma membrane blebs containing mitochondria (G). The quantification of intracellular mitochondrial “vesicles” was done from CTRL (n=21 cells), CTRL+γ-sec inh (n=26 cells), CTRL+Baf A1 (n=57 cells), APPswe (n=32 cells), APPswe+γ-sec inh (n=30 cells) and APPswe+Baf A1 (n=71 cells) (G). The quantification of plasma membrane blebs containing mitochondria was done from CTRL (n=28 cells), CTRL+γ-sec inh (n=52 cells), CTRL+Baf A1 (n=33 cells), APPswe (n=48 cells), APPswe+γ-sec inh (n=79 cells) and APPswe+Baf A1 (n=36 cells) (G). **H)** NTA quantification of the concentration of secreted EVMs obtained from CTRL (n=15), CTRL+γ-sec inh (n=13), CTRL+Baf A1 (n=4), APPswe (n=15), APPswe+γ-sec inh (n=11) and APPswe+Baf A1 (n=4) individual preparations. Scale bars= 10µm. Graphs represents means ± SEM of the indicated number of cells and from n=6 (B), n=4 (C) or (n=3) (F, G) independent experiments. (B, C, F-H) Statistical significance was determined by Kruskal Wallis test. *p value <0,05; **p value <0,01; ***p value <0,001; ****p value <0,0001; ns: not significant.

Altogether, these results demonstrate that in our study model, EVMs originate mainly from the budding of the plasma membrane and that the accumulation of APP-CTFs greatly affect the plasma membrane blebbing and secretion of EVMs.

### EVMs present a plasma membrane lipidomic profile with altered signature in APPswe EVMs

Lipid composition is crucial for EVs biogenesis, stability, and function, as well as for their interactions with target cells^33^. Previous studies have shown that while exosomes are predominantly enriched in sphingomyelin, cholesterol, and glycosphingolipids, reflecting their origin from the endosomal pathway, microvesicles retain a lipid composition similar to that of the plasma membrane.

In line with these findings, quantitative lipidomic analysis revealed that CTRL-derived exosomes and EVMs form two distinct PCA clusters (supplementary Fig. 5A). These analyses demonstrated that phospholipids (PLs) are the dominant lipids in exosomes and EVMs, with phosphatidylcholine (PC) accounting for up to 80%. Meanwhile, phosphatidylethanolamine (PE), phosphatidylserine (PS), phosphatidylinositol (PI), phosphatidylglycerol (PG), and lysophosphatidylcholine (LPC) collectively constitute approximately 7–8% (supplementary Fig. 5B). Globally, we observed a significant increase in total PC levels in EVMs compared to exosomes (supplementary Fig. 5B). An in-depth comparison of PLs types revealed no changes in minor PS levels between EVMs and exosomes (supplementary Fig. 5C). However, EVMs exhibited increased levels of PE(18:0_18:1) and PC(16:1e_18:0; 16:1_18:1; and 18:1_18:1) species, alongside reductions in PE(18:1p_20:4) and PC(16:0_14:0; 16:0_16:0; and 16:0_18:1) species (supplementary Fig. 5D, E).

Additionally, the composition of sphingolipids (SPs) differed significantly between EVMs and exosomes. EVMs showed a marked decrease in sphingomyelin (SM) levels (3.9% versus 7.9% in exosomes) with a corresponding increase in ceramide (Cer) levels (3.2% versus 0.8% in exosomes) (supplementary Fig. 5B). Further examination of fatty acid molecular profiles revealed that EVMs exhibited increased levels of Cer(d18:1_16:0 and d18:1_24:1), alongside decreased levels of all detected hexosylceramides (Hex1Cer d18:1_16:0; d18:1_16:0:22; d18:1_24:0 and d18:1_24:1) (supplementary Fig. 5F). Notably, EVMs displayed an increase in SM(d18:1_16:0) compared to exosomes (supplementary Fig. 5G).

Glycerolipids (GLs), including diglycerides (DG), triglycerides (TG), and sterol lipids (SLs) composed of cholesterol esters (ChE), represent a minor lipid class in EVs and are equivalently present in EVMs and exosomes (supplementary Fig. 5B). However, detailed analyses showed a decrease in DG(18:0_20:4) and TG(16:0_16:0_18:1) species in EVMs (supplementary Fig. 5H, I). Interestingly, while free cholesterol ChE() levels were significantly increased in EVMs compared to exosomes, esterified ChE species (ChE(18:1); (18:2); and (20:4)) were reduced (supplementary Fig. 5J). These data sustain the distinct lipidomic origins of EVMs (plasma membrane) versus exosomes (endosomes).

Since lipid dysregulation has been implicated in AD^34, 35^, we investigated whether EVMs isolated from APPswe cells exhibit a distinct lipidomic signature compared to CTRL EVMs. This comparative analysis revealed that APPswe- and CTRL-derived EVMs formed two distinct PCA clusters (supplementary Fig. 6A). While total PLs percentages remained unchanged between CTRL and APPswe EVMs (supplementary Fig. 6B), detailed analyses of fatty acid composition showed in APPswe EVMs increased levels of PS(18:0_18:1) and PE(18:0_18:1; 18:0_20:4; 18:1_18:1 and 18:1p_20:4), PC(16:1_18:1; 18:0_18:1 and 18:1_18:1). Conversely, levels of PS(18:0_22:4), PE(16:0p_22:3; 16:0p_22:6 and 18:0p_22:3), and PC(16:0_16:0; 16:0e_16:0 and 16:0_18:1) were decreased in APPswe EVMs (supplementary Fig. 6C–E). Interestingly, the lipid composition of APPswe EVM membranes showed a reduction in Cer levels (-2.1%) with a concomitant increase in SM levels (+2.3%) compared to CTRL EVMs (supplementary Fig. 6B). Furthermore, the fatty acid composition of SM and Cer revealed increased levels of Cer(d18:1_18:0) and SM(d18:1_16:0), alongside decreased levels of Cer(d18:2_16:0) and SM(d36:1) in APPswe-derived EVMs compared to CTRL-derived EVMs (supplementary Fig. 6F, G). Finally, we observed a slight but non-significant increase in total DG (+0.3%) and TG (+0.8%) levels in APPswe-derived EVMs compared to CTRL EVMs (supplementary Fig. 6B). However, while DG fatty acid composition remained unchanged, we noticed a significant decrease in TG(16:0_18:1_18:1) in APPswe EVMs versus CTRL EVMs (supplementary Fig. 6H, I). Additionally, free ChE() levels were also significantly decreased in APPswe EVMs compared to CTRL EVMs (supplementary Fig. 6J).

These data support a differential lipidomic signature of APPswe EVMs versus CTRL EVMs, likely influencing their biogenesis, stability, and function.

### APPswe-derived EVMs transfer mitochondrial dysfunction

We lastly investigated the impact of APPswe-derived EVMs in the transfer of dysfunctional mitochondria to control naïve cells. In a proof-of-concept experiment (Fig. 8A), we showed that EVMs labeled with MitoTracker green are internalized and fuse with mitochondria reticulum of receiving cells expressing mitRFP probe (Fig. 8B). We assessed the impact of exogenous EVMs application on mitochondrial membrane potential of recipient naïve control cells using TMRM probe. Our experimental conditions included EVMs, exosomes as well as the control fractions ΔEVMs (depleted in EVMs but containing exosomes), and ΔΔEVs (depleted in both EVMs and exosomes) (Fig. 1B). These preparations were obtained from CTRL or APPswe cells treated or not with the γ-sec inhibitor, thus investigating the potential impact of APP-CTFs accumulation (as in Fig. 6). Confocal imaging reveals an increase in the TMRM signal in control recipient cells following the application of EVMs isolated from CTRL cells treated with γ-sec inhibitor (i.e. accumulating endogenous APP-CTFs) (Fig. 8C, D). This increase was progressively pronounced with the application of EVMs isolated from untreated APPswe cells and further amplified with EVMs from APPswe cells treated with the γ-secretase (i.e. exacerbating endogenous and overexpressed APP-CTFs accumulation) (Fig. 8C, D). We observed a gradual mitochondrial membrane hyperpolarization upon the application of ΔEVMs fractions that was less pronounced versus EVMs application and larger compared to exosomes application (Fig. 8C, D). The application of the depleted fraction ΔΔEVs almost did not modify the mitochondrial membrane potential of receiving cells (Fig. 8C, D). These data indicate that EVMs, and to a lesser extent ΔEVMs and exosome applications, trigger mitochondrial membrane hyperpolarization. Such mitochondrial membrane hyperpolarization also appears to be associated with APP-CTFs load.

**Figure 8:**
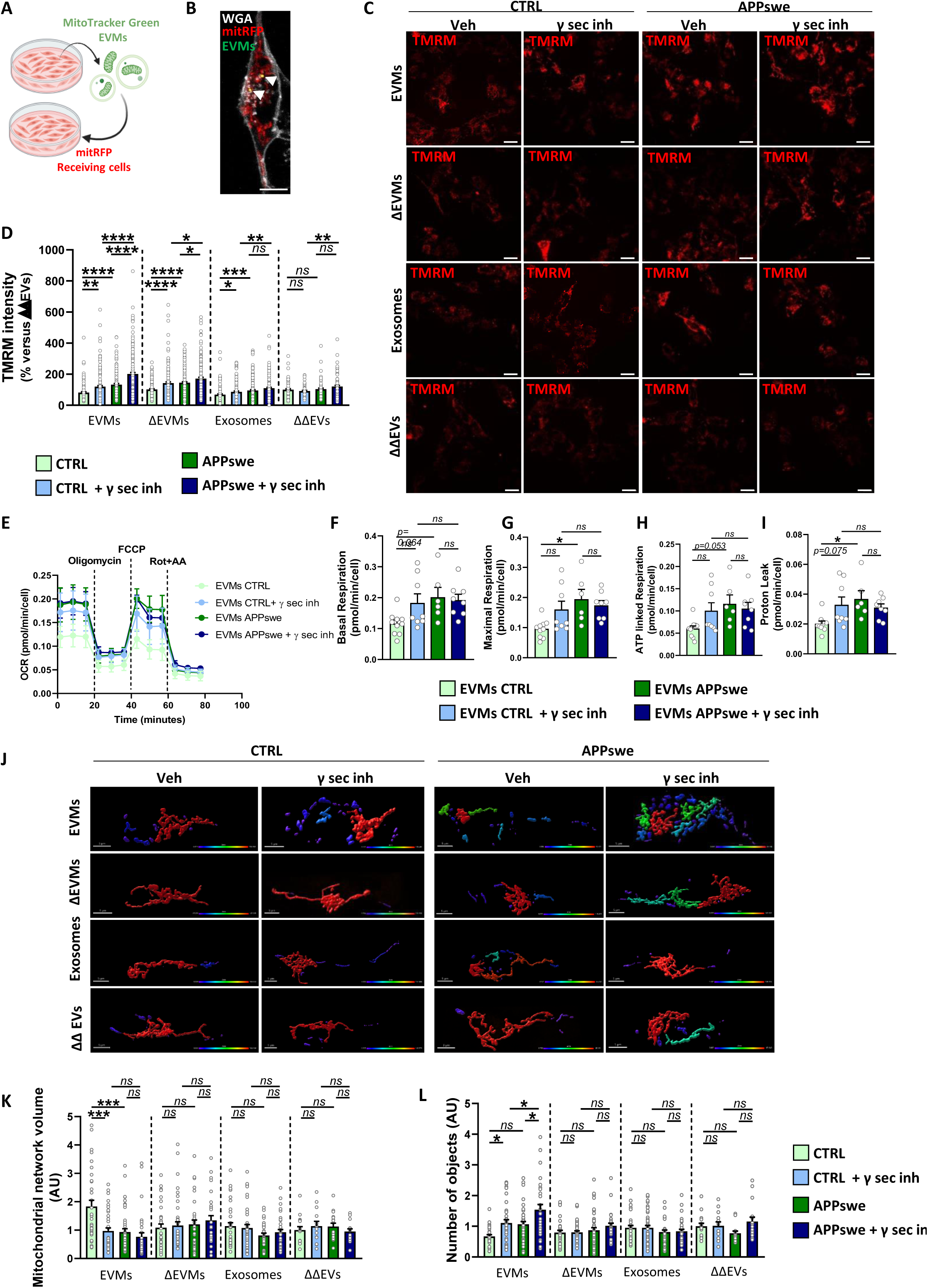
APPswe-derived EVMs spread mitochondrial dysfunction. **A)** scheme of the experimental protocol showing the transfer of isolated EVMs stained with MitoTracker Green probe and applied to receiving naïve cells transfected with mitRFP mitochondrial probe. **B)** Representative confocal image of EVMs (green) captured by a receiving cell expressing mitRFP (red) and stained with the plasma membrane dye WGA (white). **C, D)** Representative images (C) and quantitative graph (D) of TMRM mean intensity in at least n=80 receiving cells after the application of EVMs, exosomes or the control fractions either depleted in EVMs (ΔEVMs), or depleted in EVMs and exosomes (ΔΔEVs) isolated from CTRL or APPswe cells treated or not with the γ-secretase inhibitor (γ-sec inh). **E-I)** Cell respiratory capacity determined using a SeaHorse XFe96 extracellular flux analyzer (Mito Stress Test) in receiving cells treated as in (C, D). The data are expressed in pmol/min/cell ± SEM obtained from at least n=6 replicates from n=2 independent experiments. **J-L)** Representative 3D reconstruction of mitochondrial network (J) and quantitative graphs of the mitochondrial network volume (K) and the number of objects (L) in at least n=12 receiving cells treated as in (C, D). Color scale indicates large mitochondrial volume in red and small mitochondrial volume in blue. Scale bars= 10µm (B, C), or =5µm (J). Graphs represents means ± SEM of indicated n of cells. Quantified cells in (B, C and J-L) were obtained from at least (n=2) independent experiments. Statistical significance was determined by Kruskal Wallis test. *p value <0,05; **p value <0,01; ***p value <0,001; ****p value <0,0001; ns: not significant.

To further ascertain this statement, we investigated the impact of EVMs, exosomes and control depleted fractions (ΔEVMs and ΔΔEVMs) on mitochondrial respiration and measured the oxygen consumption rate (OCR) by Seahorse technology (Fig. 8E). We used a set of mitochondrial respiration modulators (oligomycin A: ATP synthase inhibitor; FCCP: mitochondrial membrane uncoupler; and rotenone/antimycin A: complex I/complex III inhibitors) to determined basal and maximal respiration (Fig. 8F, G), as well as oxygen consumption linked to ATP synthesis (Fig. 8H) and to proton leak (Fig. 8I). We reported in control naïve cells treated with APPswe EVMs, significant increase in maximal respiration (Fig. 8 G), as well as proton leak (Fig. 8I) as compared to cells that received CTRL EVMs. A trend increases of these alterations were also observed in cells receiving EVMs obtained from CTRL and APPswe cells treated with the γ-sec inhibitor (Fig. 8F-I). Importantly, the application of exosomes or ΔEVMs or ΔΔEVs fractions to control naïve receiving cells did not impact mitochondrial basal or maximal respiration (supplementary Fig. 7A, B), ATP-linked respiration (supplementary Fig. 7C) or proton leak levels (supplementary Fig. 7D).

As mitochondrial function is intimately linked to mitochondrial structure, we evaluated the impact of the exogenous application of EVMs, exosomes or ΔEVMs or ΔΔEVs fractions on mitochondrial volume network and objects number structure using 3D reconstructed images of cells expressing mitRFP probe (Fig. 8J). Control naïve cells show a gradual decrease of mitochondrial network volume when treated with EVMs isolated from CTRL cells treated with the γ-secretase inhibitor and from APPswe cells untreated or treated the γ-secretase inhibitor alongside with a gradual increase of the number of mitochondria. We further showed that exosomes, ΔEVMs, or ΔΔEVs application did not impact mitochondrial volume network or number of objects (Fig. 8J-L) as it was the case for mitochondrial respiration (supplementary Figure 7).

Overall, these data demonstrate that APPswe-derived EVMs transfer mitochondrial alterations to naïve receiving cells. We assume that these alterations are linked to dysfunctional mitochondrial content, specific proteomic and lipidomic signatures and to some extent, to APP-CTFs load.

## DISCUSSION

While small EVs (exosomes) have been extensively studied in AD^15, 16^, our study primarily focuses on the structural, molecular and functional characterization of the less well-known large extracellular vesicles containing mitochondria (EVMs) in AD cells. Specifically, we investigated the secretion of EVMs and the role of AD-derived EVMs in transferring mitochondrial pathology to healthy cells.

Previous studies have demonstrated the role of exosomes in the propagation of proteinopathies, serving as carriers of APP, APP-CTFs, and Aβ. This has been shown in exosomes isolated from cellular models of AD, including murine (N2a) and human (SH-SY5Y) neuroblastoma cells, as well as HEK293 cells expressing APPswe^14, 15, 18, 36^, as well as from AD mice models^18, 37^ and human AD brains^19^.

Consistent with these findings and as recently demonstrated^18^, we show here that exosomes derived from APPswe-expressing cells contain APP and APP-CTFs. Notably, the accumulation of APP-CTFs in exosomes is amplified upon γ-secretase inhibition. The presence of APP-derived products in exosomes is supported by the trafficking, the sorting and the proteolytic processing of APP by the β-secretase and the γ-secretase in endosomes and by the fact that exosomes originate from ILVs formed within MVBs, which are late-stage endosomes^12^. Given that APP and its derived products including APP-CTFs and Aβ have also been shown to localize in mitochondria as well as in mitochondria-associated membranes (MAMs)^8, 10, 38, 39^, we detected APP and APP-CTFs in APPswe EVMS, although to a lesser extent compared to exosomes. However, dedicated research is needed to elucidate the mechanisms governing the selective packaging of APP and APP-CTFs into EVMs. Finally, we did not detect Aβ peptides in either exosomes or EVMs, likely due to the low Aβ load under our experimental conditions.

Although small EVs have been shown to harbor mitochondrial proteins and mtDNA^40^, our results demonstrated that EVMs serve as carriers of intact mitochondria organelles, either as isolated entities or surrounded by a double membrane. Specifically, we showed that EVMs isolated from the media of APPswe-expressing cells contain mitochondria with depolarized membranes and elevated levels of mitochondrial reactive oxygen species (mitROS). Furthermore, these mitochondrial dysfunctions were exacerbated in EVMs derived from cells treated with the γ-secretase inhibitor, which is known to cause an accumulation of APP-CTFs in mitochondria, leading to mitochondrial dysfunction and impaired mitophagy^10^. These findings demonstrate that EVMs derived from AD cells carry dysfunctional mitochondria and may play a distinct role in the intracellular transport and distribution of APP metabolites, potentially contributing to the pathophysiology of AD.

In addition to carrying intact mitochondria, we show that EVMs are enriched in mitochondrial proteome compared to exosomes. This diversity in the proteomic profile further underscores the distinction between exosomes and EVMs specific biological functions. The differential proteomic analyses of APPswe EVMs versus CTRL EVMs revealed a down-regulation of proteins linked to antioxidant activity in APPswe EVMs highlighting a pathogenic signature linked to oxidative stress of AD EVMs^41^. This analysis also highlights an up-regulation of endoplasmic reticulum (ER) related functions such as unfolded protein binding or ER-associated protein degradation. Interestingly, adverse ER stress responses were described in AD brains and models^42^. Moreover, a set of ER stress responses are linked to mitochondria and MAMs molecular and structural alterations^9, 38, 43^.

We further observed a significant upregulation of mitochondrial proteins in APPswe EVMs compared to CTRL EVMs. Since the number of secreted EVMs was identical between the CTRL and APPswe conditions, and that the proteomic analysis was conducted using equal amounts of EVM particles and calibrated protein quantities, we propose that APPswe EVMs may contain a higher number of mitochondria per vesicle and/or a greater mitochondrial protein load per EVM. In support of this last statement, we reported defective mitophagy and enhanced mitochondria protein load in APPswe cells^10^. Interestingly, we similarly described dysfunctional mitochondria in sporadic AD fibroblasts^29^ and we highlight herein a specific mitochondrial proteome content in EVMs obtained from conditioned media of fibroblasts of AD patients at the prodromal stage. Consistent with our results, EVs isolated from AD patients’ brains were described to harbor a proteomic profile linked to mitochondrial metabolism as compared to control brain EVs^19^. Lipids play critical roles in EVs functions^33^. Consistent with previous studies, we report that PC represents the main PLs present in both exosomes and EVMs^44^. We also unveiled a distinct lipidomic profile enriched with Cer in EVMs and with SM in exosomes. The interplay between Cer and SM is critical for regulating the structural and functional properties of EVs. In fact, SM is the main component of lipid-rafts found on the plasma membrane and is a key actor for exosome membrane integrity and cargo sorting^35^. Finally, EVs are known to be enriched in ChE^33^. However, we revealed that EVMs are enriched with unesterified free ChE, while exosomes harbor higher levels of esterified ChE. To be noted, that ChE esterification may disrupt lipid raft dynamics, influencing APP processing and can also impair mitochondrial function^39^. Therefore, changes in EVMs lipid composition might reflect cellular mitochondrial dysfunctions. The analyses of lipids signature in APPswe versus CTRL EVMs showed similar levels of PC and PE, considered the main components of the mitochondrial membrane with cardiolipin (undetected in our analyses)^45^. The accurate analysis of fatty acid chain revealed several significant changes in plasmalogen PE (pPE) species in APPswe EVMs versus CTRL EVMs. Interestingly, plasmalogens species contain enolether on glycerol that sensitize to oxidative stress and mitochondria dysfunction^45^. Differential PE species were similarly reported in APPswe expressing cells and in post-mortem human AD brains^46^. Furthermore, we uncovered a decrease in Cer levels in favor of an increase in SM levels in APPswe EVMs versus CTRL EVMs. Interestingly, proteomic analyses also highlighted a downregulation of sphingolipid signaling pathway in APPswe EVMs. However, inconsistent data from the literature point to increased Cer levels in AD^47, 48^. Both SM and Cer are known to play a role in neuronal signaling^34^ and alterations levels of SPs are suggested to cause synaptic dysfunction and neuronal apoptosis in AD^35^. Importantly, the accumulation of C99 in MAMs increases SPs turnover, thus culminating to altered lipid composition of MAMs and to defects of mitochondrial respiration^39, 49, 50^.

Previous studies have reported that factors affecting autophagy and or lysosomal functions would increase the secretion of small EVs originating from the endosomal pathway^31^. We demonstrate herein that EVMs secretion is potentiated by APP-CTFs and unlike exosomes occurs through the budding of the plasma membrane independently from the MVB secretory pathway (i.e. CD63^+^ compartments). We may assume that EVMs secretion is specifically linked to mitochondrial and mitophagy defects linked to APP-CTFs accumulation^8, 10^. Consistently, it has been proposed that increased extracellular mitochondria release could compensates for the inadequate mitophagy capacity^51, 52^.

Small EVs containing toxic cargoes were shown to be released to migrate to recipient cells and elicit their cytotoxic effect^15^. After, demonstrating that EVMs are captured by receiving cells and fuse with mitochondria organelles in cells, we demonstrated that APPswe EVMs spread mitochondrial alterations triggering mitochondrial hyperactivity (enhanced mitochondrial potential and mitochondria respiration under stress condition) linked to mitochondria fragmentation. It was proposed that hypermetabolism is a conserved feature of mitochondrial OxPhos defects^53^. Accordingly, fibroblasts isolated from sporadic AD show some aspects of mitochondrial hyperactivity^29^ and App knock-in murine AD neurons harbor hypermetabolism^54^. Moreover, proteomic analysis of APPswe EVMs reveals an upregulation of reactive oxygen and nitrogen species (RONS) pathways, which are known to induce an energetic crisis by disrupting the mitochondrial respiratory chain and promoting mitochondrial fragmentation^55^. Interestingly, Phinney et al., demonstrated that EVs from mesenchymal stem cells containing depolarized mitochondria induced hypermetabolism in receiving macrophages^52^. It was suggested that the extracellular release of depolarized mitochondria serve as warning signals to the receiving cells which could respond with enhanced bioenergetics and mitochondrial respiration of recipient cells^56, 57^.

Overall, our study demonstrates that EVMs are not merely waste carriers but may directly contribute to cellular demise in AD through the transfer of defective mitochondrial content. This line of research holds significant promise for advancing our understanding of AD and its therapeutic landscape. Targeting pathways that regulate EVM formation, release, or uptake in recipient cells could help limit the progression of mitochondrial dysfunction in AD. Additionally, studying the proteomic and lipidomic signatures of EVMs could aid in identifying new biomarkers in peripheral biofluids of human AD subjects.

## METHODS

### Patients and control individuals

Human primary fibroblasts were obtained from the IMABio3 study cohorts (NCT01775696 and NCT02576821-EudraCT2015-000257-20) (CTRL, n = 4; AD-MCI, n = 6; AD-D, n = 4), which were approved by the French Ethics Committee. All the subjects provided written informed consent prior to participating. Controls and patients with AD were included as previously described^29^. The demographic characteristics of the patients according to diagnostic group are described in supplementary Table 1.

### Chemicals and antibodies

When indicated, cells were treated overnight (O.N) with 5 µM γ-secretase inhibitor (ELND006) (Elan Pharmaceuticals, South San Francisco, CA)^58, 59^ or with 100 nM Bafilomycin A1 (Baf A1) (CM110-0100, Enzo Life Sciences). Control cells were treated with DMSO as vehicle. Primary antibodies used in the study are listed in supplementary Table 2.

### Cell culture

Cells were cultured at 37°C in a humidified atmosphere containing 5% CO2. Human SH-SY5Y cells (CRL-2266; American Type Culture Collection, Manassas, VA) stably expressing pcDNA3.1 (CTRL) or APP protein containing the familial double Swedish mutation (K595N/M596L) (APPswe), were generated as previously described^60^ and were maintained in the presence of 450 µg/mL geneticin (TO-G048, Euromedex). SH-SY5Y neuroblastoma naïve cells were cultured in Dulbecco’s modified Eagle’s medium supplemented with 10% heat-inactivated fetal bovine serum (FBS), penicillin (100 U/ml) and streptomycin (50 μg/mL).

Primary fibroblasts were cultured in DMEM (Dulbecco modified Eagle’s minimal essential medium) supplemented with 10% FBS, 1% pyruvate sodium, Penicillin (100 U/mL) and streptomycin (50 μg/mL). We used conditioned media of fibroblasts grown between the 3^rd^ and the 12^th^ passages.

### Mitochondrial fractionation

Mitochondrial crude fractionation was obtained as already described^10^. Briefly, cells were harvested with 0,05% trypsin (25300-054, Gibco), washed and resuspended in isolation buffer (250 mM D-Mannitol, 5 mM HEPES pH 7.4, 0.5 mM EGTA, and 0.1% BSA) supplemented with protease inhibitor mixture (P2714-1BTL, Sigma). After mechanical disruption using a glass Dounce homogenizer, the lysates were centrifuged at 2500 × g at 4°C for 5 min. The supernatant was then centrifuged at 10,000 × g at 4°C for 10 min to pellet mitochondrial fraction.

### Extracellular vesicles isolation

The experimental procedure is summarized in Figure 1A. The first day, cells from each cell line were seeded in six 150mm culture dishes per condition in serum containing media. The medium was changed to serum-free conditioned media (CM) and collected 24 hours later to isolate EVMs and exosomes (from approximatively 50 000 000 cells). The CM were first spun at 400 x g for 5 min to remove dead cells. The supernatant was spun again at 1500 x g for 10 min to remove cell debris. EVMs were pelleted by ultracentrifugation of the pre-cleared CM at 18,000 x g for 40 min at 4°C (type 45Ti rotor, Beckman Coulter Optima L90K Ultracentrifuge). After a washing step in PBS, the pellet of EVMs was resuspended in PBS. Exosomes were recovered from the resulting supernatant following a filtration step (0,2 µm filter) and ultracentrifugation at 100,000 x g for 2 hours at 4°C, followed by a washing step in PBS. Finally, the exosomes pellet was resuspended in PBS. We used EVMs-depleted supernatant (ΔEVMs) and exosomes-depleted supernatant (ΔΔEVs) as control fractions.

### Nanoparticles Tracking Analysis (NTA)

Both particles size and concentration were determined by nanoparticles tracking analysis (NTA) using the ZetaView system (Particle Metrics, Meerbusch, Germany) (ZetaView Software 8.02.31). Polystyrene beads (100 nm) were used to calibrate the instrument prior to sample readings following the manufacturer’s instructions. Each pellet was diluted in particle-free, 0,2 µm-filtered PBS to reach the appropriate concentration of particles (1x10^8^ to 1x10^12^ particles/ml) (between 100 and 200 particles per field of view). Brownian motion was captured by performing 11 different cell positions with repeated 30 frame rate per second video recordings. Exosomes parameters set were as follow: sensitivity: 80; shutter speed: 100; minimum brightness: 25; minimum area: 5 pixels; maximum area: 1000 pixels. EVMs parameters set were as follow: sensitivity: 80; shutter: 100; minimum brightness: 30; minimum area: 10 pixels; maximum area: 1000 pixels.

### Transmission electron microscopy

Pellets of EVMs were fixed in 1.6% glutaraldehyde in 0.1 M phosphate buffer (pH 7.4), rinsed with cacodylate buffer 0.1 M, and then post-fixed in osmium tetroxide (1 % in cacodylate buffer) reduced with potassium ferrycyanide (1 %) for 1 hour. Samples were dehydrated with several incubations in increasing concentrations of ethanol or acetone, respectively, and embedded in epoxy resin (EPON), and 70 nm ultrathin sections were contrasted with uranyl acetate and lead citrate and observed with a Transmission Electron Microscope (JEOL JEM 1400) operating at 100 kV and equipped with a Olympus SIS MORADA camera.

### SDS-Page analysis

Each pellet of EVMs or exosomes isolated from approximatively 50 000 000 cells were resuspended in a same volume of RIPA buffer (Tris 50 mM; pH 7.4 containing NaCl (150 mM), EDTA (1 mM), Triton X100 (1 %), deoxycholate (0.5 %), SDS (0.1 %) and protease inhibitor cocktail (Sigma). This allow us to quantify EVs proteins, independently of the number of vesicles. While EVMs proteins were prepared in 5X Laemmli buffer and then boiled (3min, 95°C), exosome proteins were similarly prepared in 5X Laemmli buffer but not boiled. For crude mitochondrial fraction, 3 µg of protein were loaded. EVMs, exosomes and mitochondrial fraction were separated on 16% Tris-Tricine gels to detect full length-APP, C83 and C99, or on Bio-Rad stain-free^TM^ 12% TGX FastCast^TM^ acrylamide gels (Bio-Rad, France) to detect CD63, Alix, Flotilin-2, CD81, HSP60, TOMM20, TIMM23, and HSP10. After migration, Bio-Rad stain-free^TM^ 12% TGX FastCast^TM^ acrylamide gels were photoactivated using a ChemiDoc^TM^ Touch Imaging System (Bio-Rad, France) for the visualization of total proteins (« protein stain ») before being electrophoretically transferred to nitrocellulose membranes using the Bio-Rad Trans-Blot® Turbo^TM^ Transfer system. Membranes were blocked with 5% non-fat milk in TBS-T for 1 hour at room temperature (RT), then incubated with primary antibodies O.N at 4°C. Goat anti-mouse or anti-rabbit HRP conjugated secondary antibodies (115-036-003 and 111-036-045 respectively, Jackson Immuno Research) were then used. Images were captured with ImageQuant LAS4000 using WesternBright^TM^ Sirius^TM^ HRP substrate (Advansta, France). Peak height of signal intensities from protein bands were quantified with ImageJ/Fiji software.

### Flow cytometry analyses

EVMs and mitochondrial fraction were resuspended with PBS, and stained with 100 nM MitoTracker Green^TM^ FM (MitoT) (M7514, ThermoFisher Scientific), with 2 nM TMRM probe (87919, Fluka Analytical) or with 5 µM MitoSOX^TM^ red mitochondrial superoxide indicator (M36008, ThermoFisher Scientific) at 37°C for 30 min. Samples were stained with Annexin V (BD 550474, Fisher Scientific) in 1X Annexin Binding Buffer (V13246, ThermoFisher scientific) diluted at 1/20 for 15 min at RT applied after MitoTracker Green staining. Data were acquired with Novocyte Flow Cytometer using the NovoExpress software (Agilent). The gating excluded residual debris and doublets. Single stained and unstained controls were also applied for compensations. The results represent the median intensity values for MitoTracker Green, TMRM or MitoSOX.

### Lipidomic analyses

Lipids from exosomes or EVMs were extracted according to a modified Bligh and Dyer protocol^61^ with a mixture of methanol and chloroform (2:1). Next, the mixtures were processed as described elsewhere^62^. Data were reprocessed using Lipid Search 4.2.21 (ThermoFisher). The product search mode was used and the identification was based on the accurate mass of precursor ions and MS2 spectral pattern. The parameters for the ion identification and alignment were set up as follow: ion precursor tolerance 5 ppm; ion product tolerance 5 ppm; ion and adducts searched [H+], [NH4+] and [Na+]; alignment retention time tolerance 0.4 min; ID quality filters A, B and C.

### Proteomic analyses

In-gel digestion: EVMs and exosomes pellets were prepared as described above (i.e. SDS-Page analysis section) and loaded on 12% TGX™ precast polyacrylamide gels (4561043, Bio-Rad, France). Following migration, gels were stained with silver nitrate (SilverQuest^TM^ LC6070, ThermoFisher Scientific) for the accurate calibration of proteins load. We adjusted the deposit volume according to the sample with the lowest protein content corresponding to 1x10^8^ particles. EVMs and exosomes were loaded on a second precast gel (4561094, Bio-Rad, France), which was stained with Coomassie blue (24615, ThermoFisher Scientific) for 2 hours followed by 3 washes in pure water, the last wash was O.N. Protein spots were manually excised from the gel and processed as previously described^62^. Briefly, gel pieces were reduced with 10 mM dithiothreitol (DTT) and alkylated with 55mM iodoacetamide (IAA). Gel pieces were washed twice with H2O/acetonitrile (ACN) (1/1) and once with ACN. Next, gel pieces were digested using 50 mM NH4HCO3 buffer containing 10 ng/mL of trypsin. Tryptic peptides were isolated by extraction with (i) 1% formic acid (FA) in water and (ii) ACN. Peptide extracts were pooled, concentrated under vacuum and solubilized in 15 µL of aqueous 0.1% FA and then injected.

NanoHPLC-HRMS analysis: NanoHPLC-HRMS analysis was performed using a nanoRSLC system (ultimate 3000, Thermo Fisher Scientific) coupled to an Easy Exploris 480 (Thermo Fisher Scientific) as previously described^58, 59^. Five microliter of peptides solution were injected and concentrated for peptide separation using the Easy-nLC ultra high-performance LC system. Next, peptides separation was performed on a 75 mm i.d. x 500 mm (2µm, 100 Å) PepMap RSLC C18 column (ThermoFisher Scientific) at a flow rate of 300 nL/min. Solvent systems were: (A) 100% water, 0.1%FA, (B) 100% ACN, 0.1% FA. The following gradient was used t = 0 min 2% B; t = 3 min 2 % B; t = 103 min, 20 % B; t = 123 min, 32 % B; t = 125 min 90 % B; t = 130 min 90 % B; (temperature set at 40°C). MS and MS/MS spectra were acquired as described elsewhere^63^.

MS data reprocessing: Qualitative data analysis were done using Proteome Discoverer v2.5 equiped with Sequest HT. At least 2 unique peptides per protein were required for a protein identification. For Label-Free Quantification (LFQ) analysis, MS raw files were processed using PEAKS Online X (build 1.8, Bioinformatics Solutions Inc.). Data were searched against the SwissProt Homo Sapiens database (01_2023, total entries 20405). A mass accuracy of 10 ppm was used to precursor ions with fragment mass tolerance of 0.02 Da for-product ions. Specific tryptic cleavage was selected with a maximum of 2 missed cleavages allowed. For identification, the following post-translational modifications were included: Acetylation (Protein N-term), Oxidation (M), Deamidation (NQ) as variables and carbamidomethylation (C) as fixed. Identifications were filtered based on a 1% FDR (False Discovery Rate) threshold at both peptide and protein group levels. Label free quantification was performed using the PEAKS Online X quantification module, allowing a mass tolerance of 10 ppm, a CCS error tolerance of 0.02 and a one min retention time shift tolerance for match between runs. Protein abundance was inferred using the top N peptide method and TIC was used for normalization. Multivariate statistics on proteins were performed using Qlucore Omics Explorer 3.8 (Qlucore AB, Lund, SWEDEN). The transformed data were finally used for statistical analyses using a Student’s bilateral t-test and assuming equal variance between groups. A p-value ≥0.01 was used to filter differential protein expression in APPswe EVMs versus CTRL EVMs and a p-value ≥0.05 was used to filter differential protein expression in fibroblasts-derived EVMs.

Venn diagrams were generated using the open-access website Interactive Venn (https://www.interactivenn.net). Proteins were compared to the database of human mitochondrial proteins (Human MitoCarta 3.0 dataset) (https://www.broadinstitute.org). Raw data and lists of proteins obtained after processing for heatmaps and volcano plots are supplied as supplemental material.

### Gene Ontology and bioinformatics

KEGG and Gene Ontology (GO) analysis were done using the Database for Annotation, Visualization and Integrated Discovery (DAVID) knowledgebase (v2023q2). The top 10 hits from KEGG pathways and GO cellular component (CC), molecular function (MF) and biological process (BP) were plotted according to –log10 (p value). Protein-protein interaction network analyses were done using open-access website STRING (https://string-db.org). For this analysis, an interaction score of 0,7 (high confidence based on default active interaction sources) was set as minimum and GO enrichment data were used to manually cluster proteins based on mitochondrial BP, providing a detailed map of mitochondrial proteome of EVMs.

### Colocalization of mitochondria with the plasma membrane or CD63 positive vacuoles

CTRL or APPswe cells seeded on glass coverslips were transiently transfected the mitochondrial Mit-GFP probe using Lipofectamine 2000 (11668019, Thermofisher Scientific) according to the manufacturer’s instructions. Transfected cells were treated 24 hours later with γ-secretase inhibitor or Bfa A1, and fixed with 4 % paraformaldehyde for 10 min at RT. Cells were washed three times with 1X PBS, and then permeabilized with 0.3 % Triton X-100 for 5 min. Non-specific binding sites were blocked for 1 hour in 3 % PBS-BSA containing 0.05 % tween at RT. Primary antibody for CD63 was diluted in PBS-BSA 0.3 % containing 0.005 % tween and applied O.N. After three washes with 1X PBS, AlexaFluor-conjugated secondary antibody (Jackson ImmunoResearch) was applied for 1 hour. Nuclei were stained with DAPI (1/10000, D1306, Invitrogen) and coverslips were mounted on glass slides with Vectamount medium (H-5501-60, Vector).

Live CTRL and APPswe cells were stained with the mitochondrial probe MitoTracker Green (M7514, ThermoFisher Scientific) for 30 min at 37°C, washed with PBS 1x and stained with the plasma membrane probe Wheat Germ Agglutinin (WGA, W32466, ThermoFisher Scientific) for 10 min at RT. Live cell imaging was done in KRB buffer (135 mM NaCl, 5 mM KCl, 1 mM MgSO4, 0.4 mM K2HPO4, 20 mM HEPES, 0.05 mM CaCl2, 1g/L Glucose, pH 7.4). Confocal images were acquired using Leica TCS SP5 microscope (for fixed cells) or LSM 780 (for live cells) with appropriate 63X/1.4 NA oil objective. The quantification of the colocalization (Mander’s or Pearson’s coefficient) were determined using the Fiji plug-in JACoP (Just Another Colocalization Plug-in) (https://imagej.net/contribute/citing)^64^. The quantification of the number of intracellular vesicles or plasma membrane blebs containing mitochondria per cell were done manually. Profile plots of WGA and MitoTracker Green stainings were obtained after the projection of 4 z-stacks of each image using Fiji.

### Quantification of the mitochondrial volume and of membrane potential in receiving cells

Naïve neuroblastoma cells were seeded on chambered coverglass (4 wells-CellView, 627870, Dutcher (200 000 cells/well)), and transfected twenty-four hours later with the mitochondrial Mit-RFP probe using Lipofectamine 2000 (11668019, Thermofisher Scientific) according to the manufacturer’s instructions. Twenty-four hours later, we applied equal EVMs or exosomes particles (1x10^8^ particles) or equal volumes of the respective depleted fractions (ΔEVMs or ΔΔEVs) isolated from CTRL or APPswe cells treated or not with the γ-secretase inhibitor. Cells were fixed 16h later with 4 % PFA for 10 min at RT and stained with DAPI, before being mounted on slides. Confocal images were taken with Leica TCS SP5 microscope with 63X/1.4 NA oil objective. The quantification of the mitochondrial volume and the number of objects was carried out thanks to 3D object counter plugin of Fiji. The mitochondrial volume was reconstructed using Imaris software (Imaris x64 v.9.6.1).

Naïve neuroblastoma cells were seeded as described above and stained with the Tetramethyl rhodamine methyl ester (TMRM) (87919, Fluka Analytical) at 2 nM for 30 min, at 37°C. Images with z-stacks were acquired by live imaging using Zeiss LSM 780 with 63X Objective. TMRM fluorescence median intensities was analyzed after the maximum intensity projection of z-stacks using Fiji software.

### SeaHorse analyses

Measurements of aerobic respiration were conducted as previously described^29^ with the Seahorse Bioscience XFe96 bioanalyzer (XF Mito Stress Test Kit, 103015-100, Agilent). Briefly, we seeded 50 000 cells per well on XFe96 cell culture microplates (102416-100, Agilent) two days before the experiment. One day before the experiment, 1x10^8^ particles (exosomes or EVMs) or equal volumes of respective depleted fractions (ΔEVMs or ΔΔEVs) were applied in each well O.N. On the day of the experiment, the culture medium was replaced with Seahorse Base Medium (103334-100, Agilent) supplemented with 1 mM pyruvate, 2 mM glutamine and 10 mM glucose, and incubated for 30 min at 37°C in a CO_2_-free incubator. After measuring basal respiration, drugs were added in each well in a sequential order (5 µM oligomycin, 2 µM FCCP, and 0.5 µM rotenone/antimycin A). Cells were fixed with PFA 4 % and stained with DAPI diluted at 1/10 000 (Roche) for 5 min at RT. Cytation 5 Biotek was then used to count the cell nuclei and normalize the experiments on the relative cell number per well. Data were analyzed using the XF Cell Mito Stress Test Report Generator.

### Statistical analyses

Data were expressed as means ± SEMs. Statistical analyses were carried out with GraphPad Prism version 9 for Windows (GraphPad Software, La Jolla, CA, USA; https://www.graphpad.com). The sample size and statistical information for each experiment are detailed in figure legends. The data were first analyzed for a normal distribution. The Mann-Whitney test was used when groups of two variables did not pass the normality test. Groups of more than two variables that have passed normality test were analyzed by a Two-Way ANOVA test followed by a Tukey’s multiple comparison test. We used one-way ANOVA Tukey’s multiple comparisons post hoc test when groups of two or more variables passed the normality test. All p values below or equal to 0,05 were considered significant: *p < 10.05, **p < 0 0.01, ***p < 0.001, ****p < 0.0001 and ns: not significant. Statistical analyses were reported between individual cells obtained from independent experiments (Fig. 7A, B and E-G; Suppl. Fig. 6 and Fig. 8C, D and J-L). We also reported in Figure legends the biological effects (Hedges test) using a free online effect size calculator (https://camel.psyc.vt.edu/models/stats/effect_size.shtml).

Microscopy images were assembled using OMERO.figure (University of Côte d’Azur and EMBRC-France). Schematic representations are created using Biorender.com.

### Data availability

Proteomic and lipidomic datasets are available upon demand.

## Acknowledgements

We wish to thank the Conseil Départemental D06 for the funding of the ZetaView and the SEAHORSE. This work was supported by Fondation Vaincre Alzheimer-Grant FVA #18035, France Alzheimer AAP PFA 2021-Grant # 6174 and by the French government, managed by the Agence Nationale de la Recherche under the Plan d’investissement France 2030, as part of the Initiative d’Excellence d’Université Côte d’Azur bearing the reference ANR-15-IDEX-01 to MC. The post-doctoral fellow salary was supported by IDEX UCAJedi AAP Jeunes Chercheurs 2022 to MC and FE, and the LabEx (excellence laboratory) DISTALZ (development of Innovative Strategies for a Transdisciplinary approach to Alzheimer’s disease) to FC. The authors acknowledge the Electron Microscopy facility CCMA (Centre Commun de Microscopie Appliquée) from the « Université Côte d’Azur », part of the « Microscopie Imagerie Côte d’Azur » GIS IBiSA labeled platform, supported by Université Côte d’Azur, the “Région Sud and the Conseil Départemental 06 and Christelle Boscagli for electron microscopy acquisitions. This study was partially funded by Institut de Recherches Servier and supported by Investissements d’Avenir grants ANR-10-IAIHU-06. We thank the DNA & Cell Bank core facilities of ICM as well as the French Health Ministry (PHRC) under reference PHRC-0054-N 2010 and PHRC-2013-0919, the Institut Roche de Recherche et Médecine Translationelle as well as CEA, the Fondation pour la recherche sur Alzheimer and France-Alzheimer to MS and MCP. The authors acknowledge Guillaume Dorothée and Michel Bottlaender for their contribution to patient data collection and Lucile Fleuriot for lipidomic analyses.

## DISCLOSURE OF INTEREST

The authors report no conflict of interest

### Author contributions

MC conceptualized the project and funded the project. MC and FE designed the experimental plans, discussed data, and wrote and revised the manuscript. FE developed the project and performed all experiments, processed and analyzed all data and prepared the figures. ASG and DB performed proteomic and lipidomic analyses. ASG, VL and GC processed and analyzed proteomic data. DB processed lipidomic data. KK contributed to imagery analyzed. SLG conducted electron microscopy experiments. MCP and MS collected patient data and provided primary cultures of fibroblasts. FC contributed to the funding of the project. All authors have read edited and agreed to the published version of the manuscript.

### Research ethics

The use of patients and control individuals-derived primary fibroblasts was approved by a French Ethics Committee. All subjects provided written informed consent prior to participating.

### Additional information Flux cytometry

EVs or mitochondrial fraction were treated with 0.5% Triton X-100 (T8787, Sigma-Aldrich) for 20 min at room temperature (RT).

## Supplementary Figures and legends

**Suppl. Figure 1:** Flow cytometry analyses of EVMs. **A, B)** Plots of additional controls representing the background of microparticles found in PBS (A) and in PBS + MitoTracker green (MitoT) (B). **C, D)** Plots of CTRL and APPswe mitochondrial fractions unstained (C) or stained with the MitoT (D). **E, F)** Plots of CTRL and APPswe mitochondrial fractions (E) or EVMs (F) stained with MitoT and incubated with Triton X-100 (+ MitoT + Triton). **G-N)** Plots of CTRL EVMs (G, I, K), or APPswe EVMs (H, J, L) stained with the MitoT (G, H) or with Annexin V (Ann V) (I, J) or double-stained with the MitoT and AnnV (+MitoT + Ann V) (K-N). **O)** Quantitative graphs of the percentage of unstained EVMs or stained with the MitoT or with MitoT+AnnV. Graph represent the mean ± SEM of n=3 individual preparations per condition. Statistical significance was determined by one-way ANOVA with Tukey’s test. ns: not significant.

**Suppl. Figure 2: A)** Venn diagrams representing the number of proteins identified in each of the four replicates of EVMs isolated from CTRL and APPswe CM. **B)** Venn diagram representing the proteins differentially identified in at least 3 out of 4 replicates of CTRL and APPswe EVMs. **C)** KEGG Pathway and **D-F)** Gene Ontology (GO) of the top 10 Cellular Component (CC) (D), Molecular Function (MF) (E) and Biological Process (BP) (F) unique to CTRL EVMs or APPswe EVMs or identified in both CTRL and APPswe EVMs. Abbreviations: LTP: long-term potentiation; LTD: long-term depression; ECM: extracellular matrix; Ser: serine; Val: valine; Leu: leucine; Ile: isoleucine; Mito: mitochondria; ER: endoplasmic reticulum; ALS: amyotrophic lateral sclerosis. The statistical significances correspond to log-transformed p-values (-log_10_ (p)).

**Suppl. Figure 3: A-C):** Venn diagrams representing the number of proteins identified in of CTRL (n=4) (A), AD-MCI (n=6) (B) and AD-D (=n=4) (C) fibroblasts-derived EVMs. **D)** KEGG Pathway and **E-G)** Gene Ontology (GO) terms of the top 10 Biological Process (BP) (E), Cellular Component (CC) (F) and Molecular Function (MF) (G). The statistical significances correspond to log-transformed p-values (-log_10_ (p)). **H-J)** Heatmap representation of the significant down-regulated (blue) and up-regulated (red) proteins in AD-D EVMs versus CTRL EVMs (H), AD-MCI EVMs versus CTRL EVMs (I) and AD-MCI versus AD-D EVMs (J).

**Suppl. Figure 4: A)** Representative images of CTRL and APPswe cells treated or not with the γ-secretase inhibitor (γ-sec inh) or with Baf A1 and stained with APP-Cter (red) and CD63 (green). Nuclei are stained with dapi (blue). The merged signal (yellow) show the colocalization of green and red signals. Scale bars = 10 μm. **B)** Pearson’s coefficient of APP C-terminal (APP-Cter) colocalization with CD63. Graph represent the mean ± SEM from CTRL (n=40), CTRL+γ-sec inh (n=36), CTRL+Baf A1 (n=43), APPswe (n=42), APPswe+γ-sec inh (n=42) and APPswe+Baf A1 (n=44). Quantified cells were obtained from n=4 independent experiments. Statistical significance was determined by Two-way ANOVA with Tukey’s test. **p value < 0,01; ***p value < 0,001; ****p value <0,0001; ns: not significant.

**Suppl. Figure 5: Comparative lipidomic analysis of exosomes and EVMs. A)** Principal component analysis (PCA) of four individual Exosomes and EVMs preparations. **B)** Composition of exosomes and EVMs lipids according to lipid classes. PE: Phosphatidylethanolamine; PC: Phosphatidylcholine; PS: Phosphatidylserine; PI: Phosphatidylisnositol; PG: Phosphatidylglycerol; LPC: Lysophosphatidylcholine; Cer: ceramid; SM: Sphingolmyelin; DG: Diglyceride; TG: Triglyceride; and ChE: cholesterol. **C-J)** Comparison of Phosphatidylserine (PS) (C), Phosphatidylethanolamine (PE) (D), Phosphatidylcholine (PC) (E), Ceramid (Cer) (F), Sphingomyelin (SM) (G), Diglyceride (DG) (H), Triglyceride (TG) (I) and Cholesterol (ChE) (J) of individual lipid molecules in CTRL exosomes and EVMs. B-J) Data represents the percentage of lipid class versus total lipid (B) or the abundance in percentage of lipid molecules versus total lipid class considered as 100% (D-J). Data ± SEM were obtained from n=4 independent preparations of exosomes or EVMs. Statistical significance was determined by Two-way ANOVA. *p value <0,05; **p value <0,01; ***p value <0,001; ****p value <0,0001.

**Suppl. Figure 6: Comparative lipidomic analysis of APPswe-derived EVMs versus CTRL-derived EVMs. A)** Principal component analysis (PCA) of four individual CTRL and APPswe EVMs preparations. **B)** Composition of CTRL and APPswe EVMs lipids according to lipid classes as described in suppl. Figure 5. **C-J)** Comparison of Phosphatidylserine (PS) (C), Phosphatidylethanolamine (PE) (D), Phosphatidylcholine (PC) (E), Ceramid (Cer) (F), Sphingomyelin (SM) (G), Diglyceride (DG) (H), Triglyceride (TG) (I) and Cholesterol (ChE) (J) of individual lipid molecules from CTRL EVMs and APPswe EVMs (n=4). B-J) Data represents the percentage of lipid class versus total lipid (B) or the abundance in percentage of lipid molecules versus total lipid class considered as 100% (D-J). Data ± SEM were obtained from n=4 independent preparations of exosomes or CTRL or APPswe EVMs. Statistical significance was determined by Two-way ANOVA. *p value <0,05; **p value <0,01; ***p value <0,001; ****p value <0,0001.

**Suppl. Figure 7: A)** Cell respiratory capacity in receiving cells after application of the fraction depleted in EVMs (ΔEVMs), exosomes and the fraction depleted in both EVMs and exosomes (ΔΔEVs) (isolated from CTRL or APPswe cells treated or not with the γ-secretase inhibitor) determined using a SeaHorse XFe96 extracellular flux analyzer (Mito Stress Test). The data are expressed in pmol/min/cell ± SEM obtained from at least 4 replicates in 2 independent experiments. Statistical significance was determined Two-way ANOVA with Tukey’s test for each group of vesicles or control. ns: not significant

**Supplementary Table 1:** Demographic characteristics of fibroblasts isolated from patient skin biopsy. The CDR-SOB (Clinical Dementia Rating, Scale Sum of Boxes) scale indicates the severity of dementia; no dementia (CDR = 0). The maximum score for the Mini-Mental State Examination (MMSE) is 30, with lower scores associated with greater cognitive deterioration. Abbreviations: CTRL control; AD-MCI Alzheimer’s disease patients with mild-cognitive impairment; AD-D Alzheimer’s disease patients with dementia; F female; M male; CDR-SOB clinical dementia rating sum of the boxes; PiB-GCI ^11^C-labelled Pittsburgh compound B, Global cortical index.

| Group | Gender | Age | CDR-SOB | MMSE | PiB-GCI |
| --- | --- | --- | --- | --- | --- |
| CTRL | M | 64 | 0 | 28 | 1.33 |
|  | M | 66 | 0 | 29 | 1.27 |
|  | F | 81 | 0 | 30 | 2.15 |
|  | M | 59 | 0 | 30 | 1.22 |
| AD-MCI | M | 62 | 1 | 25 | 2.36 |
|  | F | 70 | 1.5 | 22 | 3.58 |
|  | M | 64 | 3 | 21 | 2.84 |
|  | F | 77 | 0.5 | 24 | 1.58 |
|  | F | 55 | 3 | 20 | 2.33 |
|  | M | 70 | 3 | 28 | 2.89 |
| AD-D | F | 52 | 8 | 13 | 2.43 |
|  | F | 60 | 5 | 17 | 3.11 |
|  | F | 71 | 4.5 | 17 | 4.14 |
|  | F | 51 | 9 | 13 | 2.86 |

**Supplementary Table 2:**
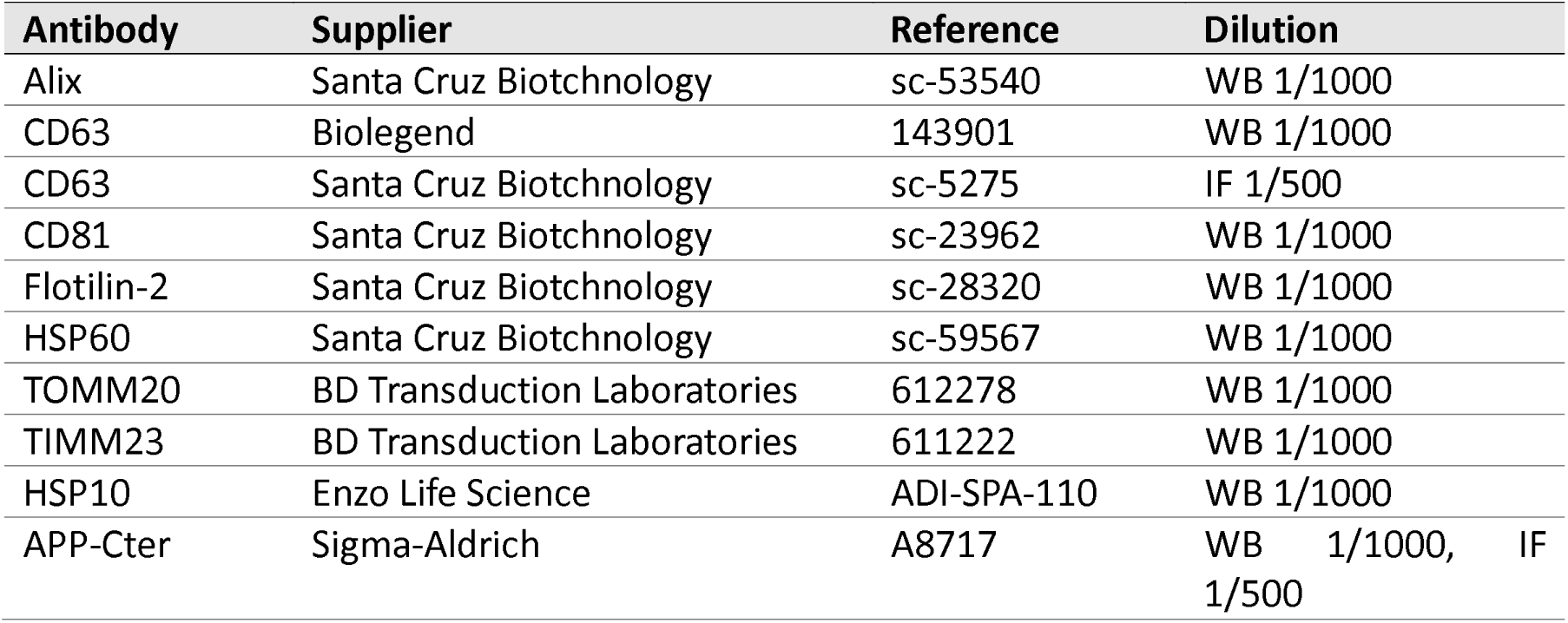
List of primary antibodies used in the study.

