## Supplementary figures for "Large extracellular vesicles containing mitochondria (EVMs) derived from Alzheimer’s disease cells harbor pathologic functional and molecular profiles and spread mitochondrial dysfunctions"

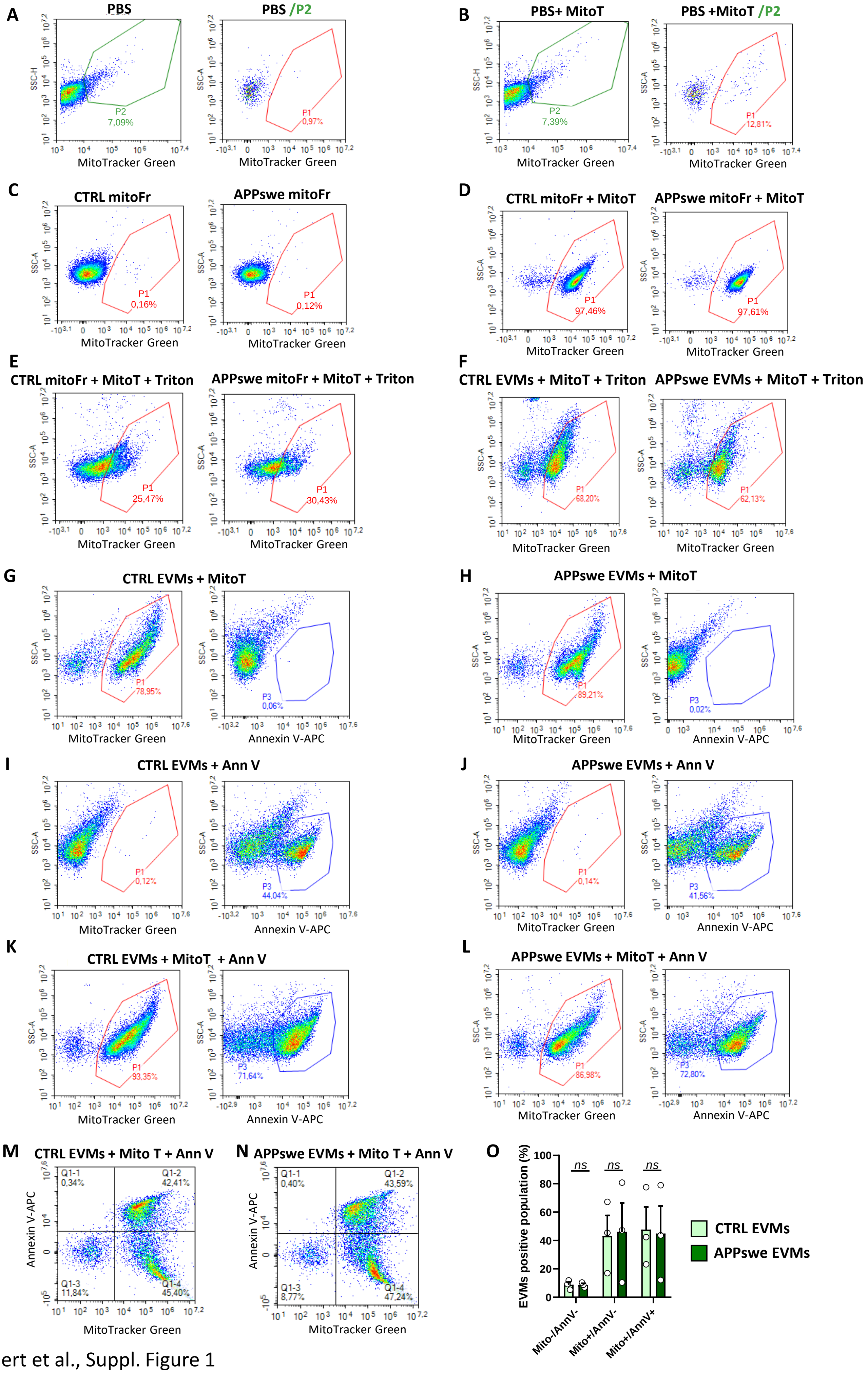


Eysert et al., Suppl. Figure 1

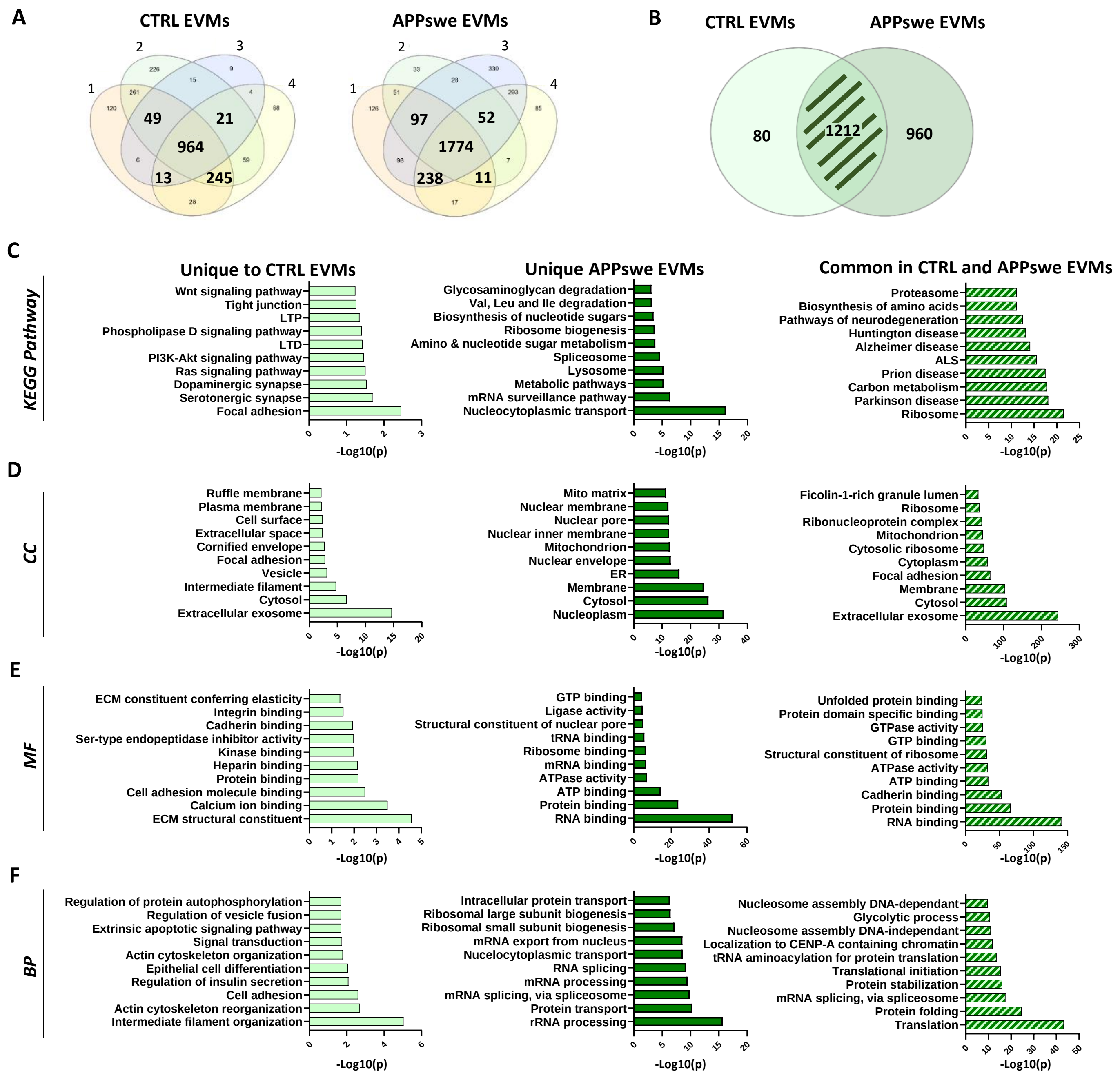


Eysert et al., Suppl. Figure 2

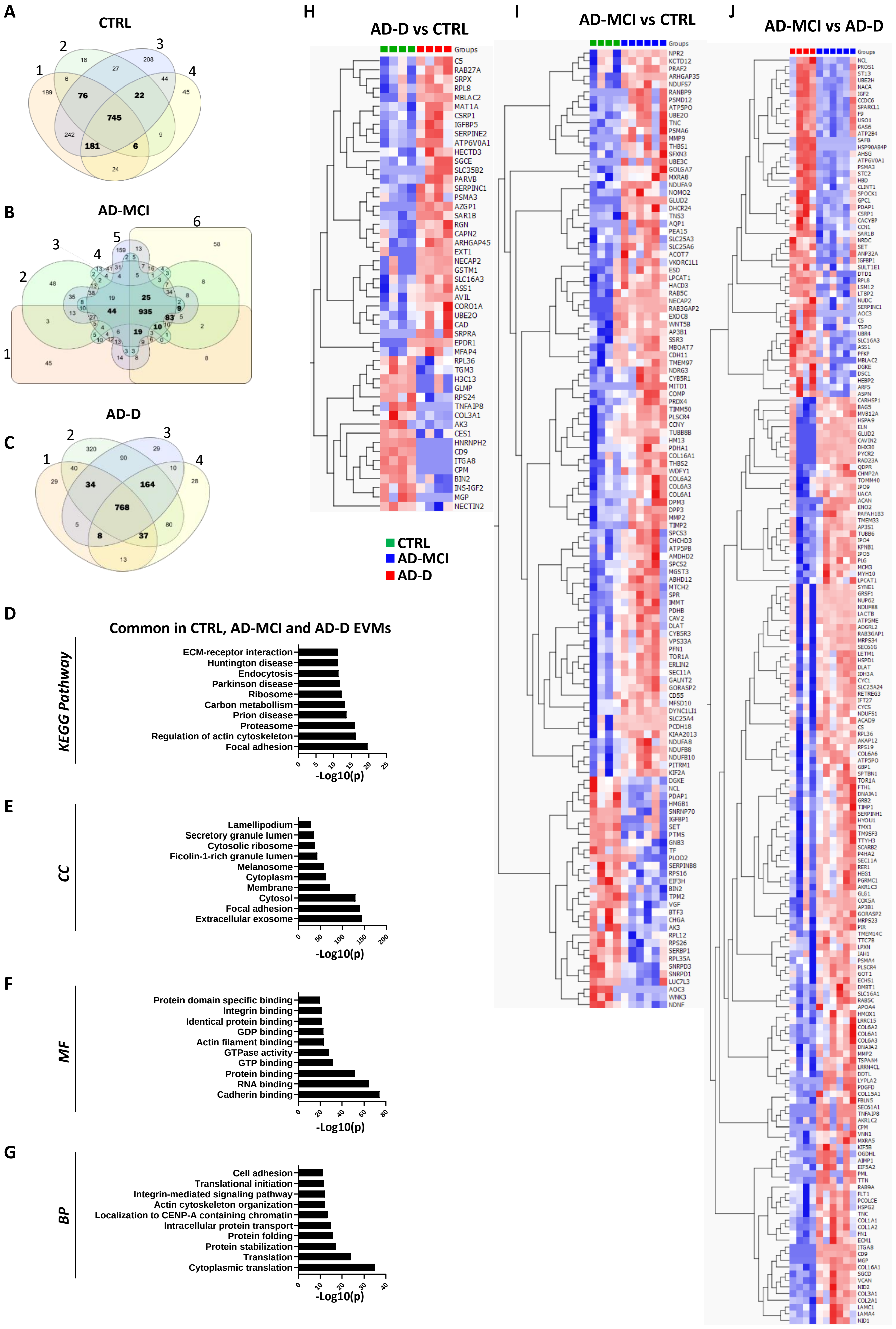


Eysert et al., Suppl Figure 3

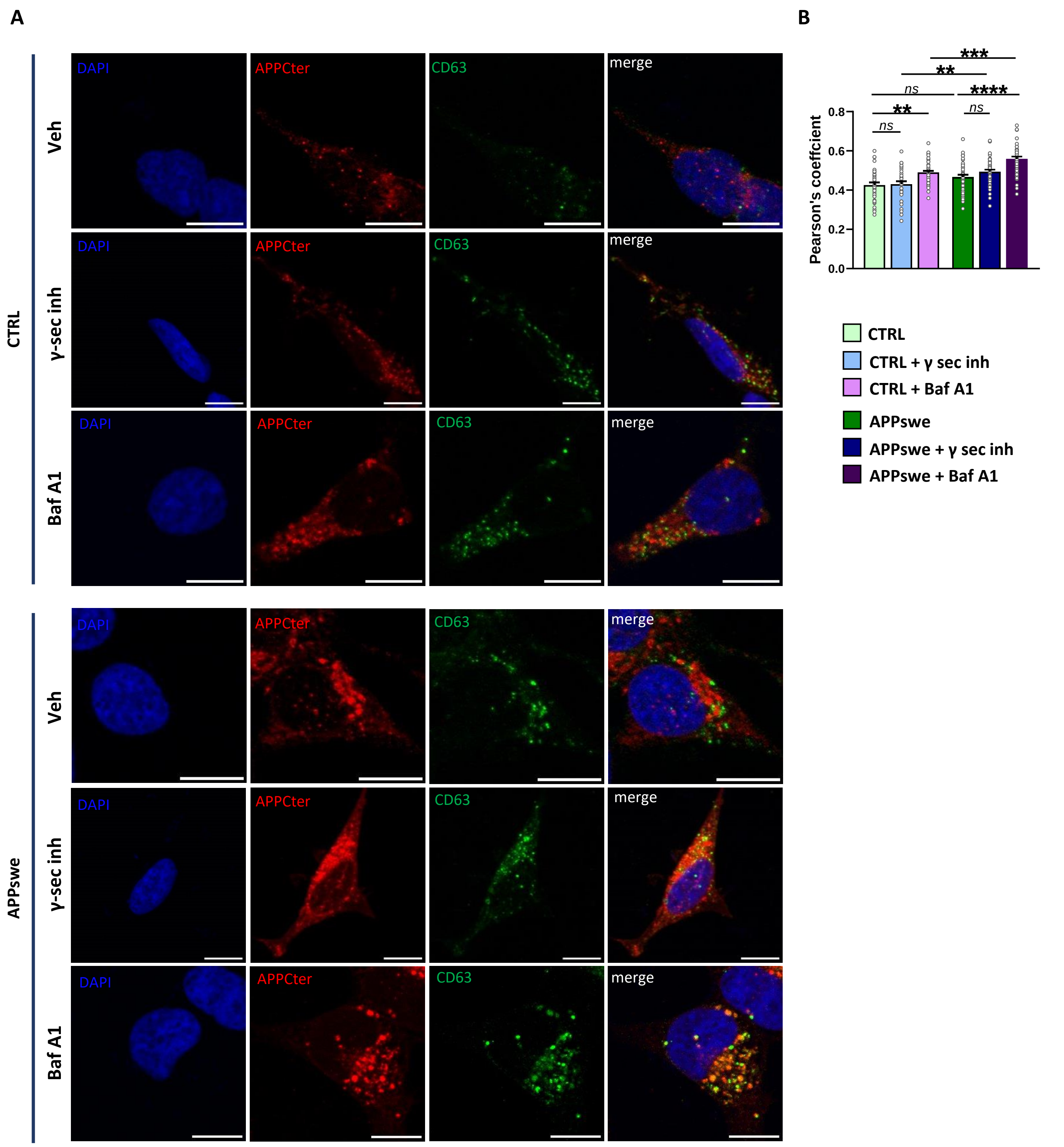


Eysert et al., Suppl. Figure 4

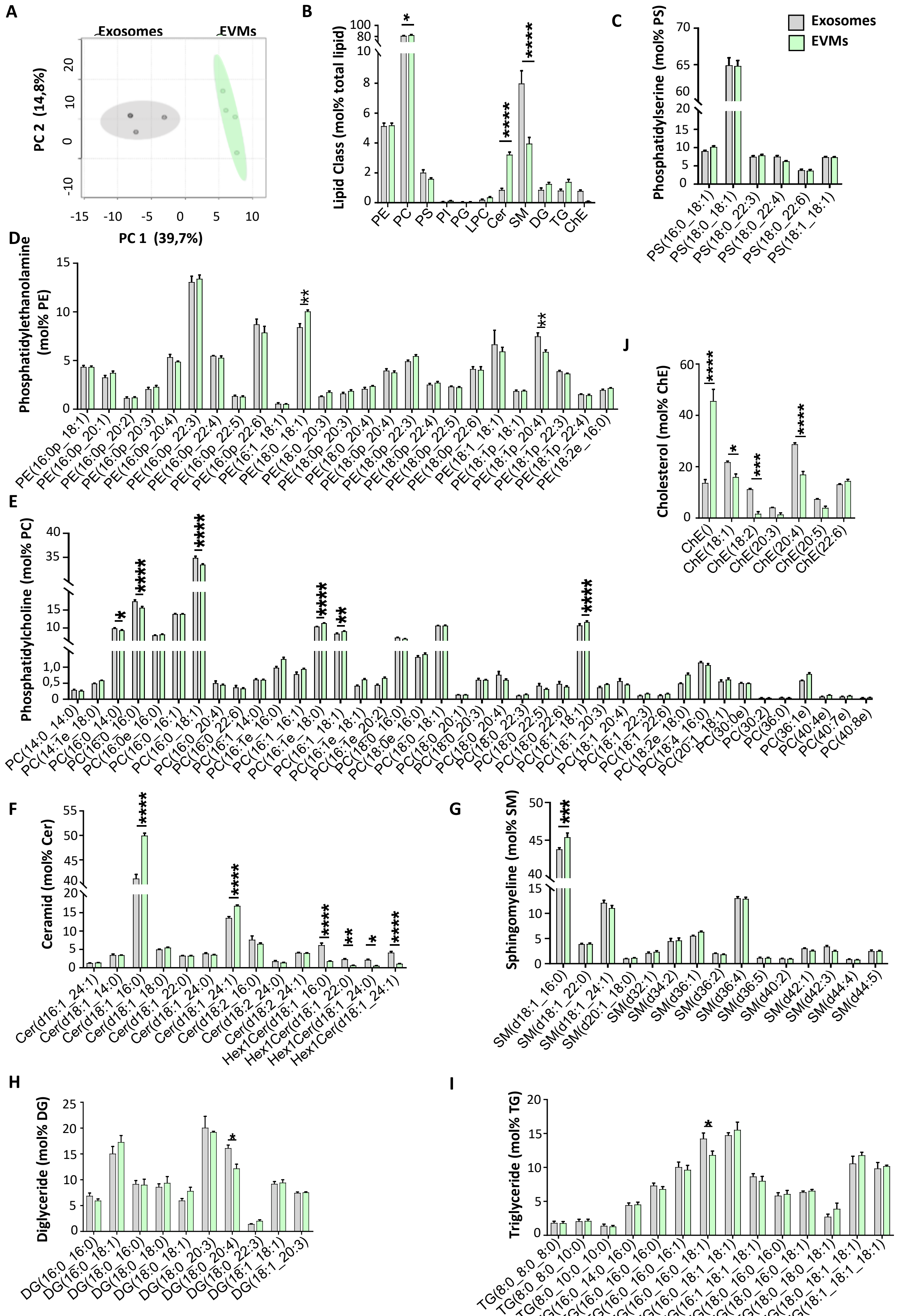


Eysert et al., Suppl. Figure 5

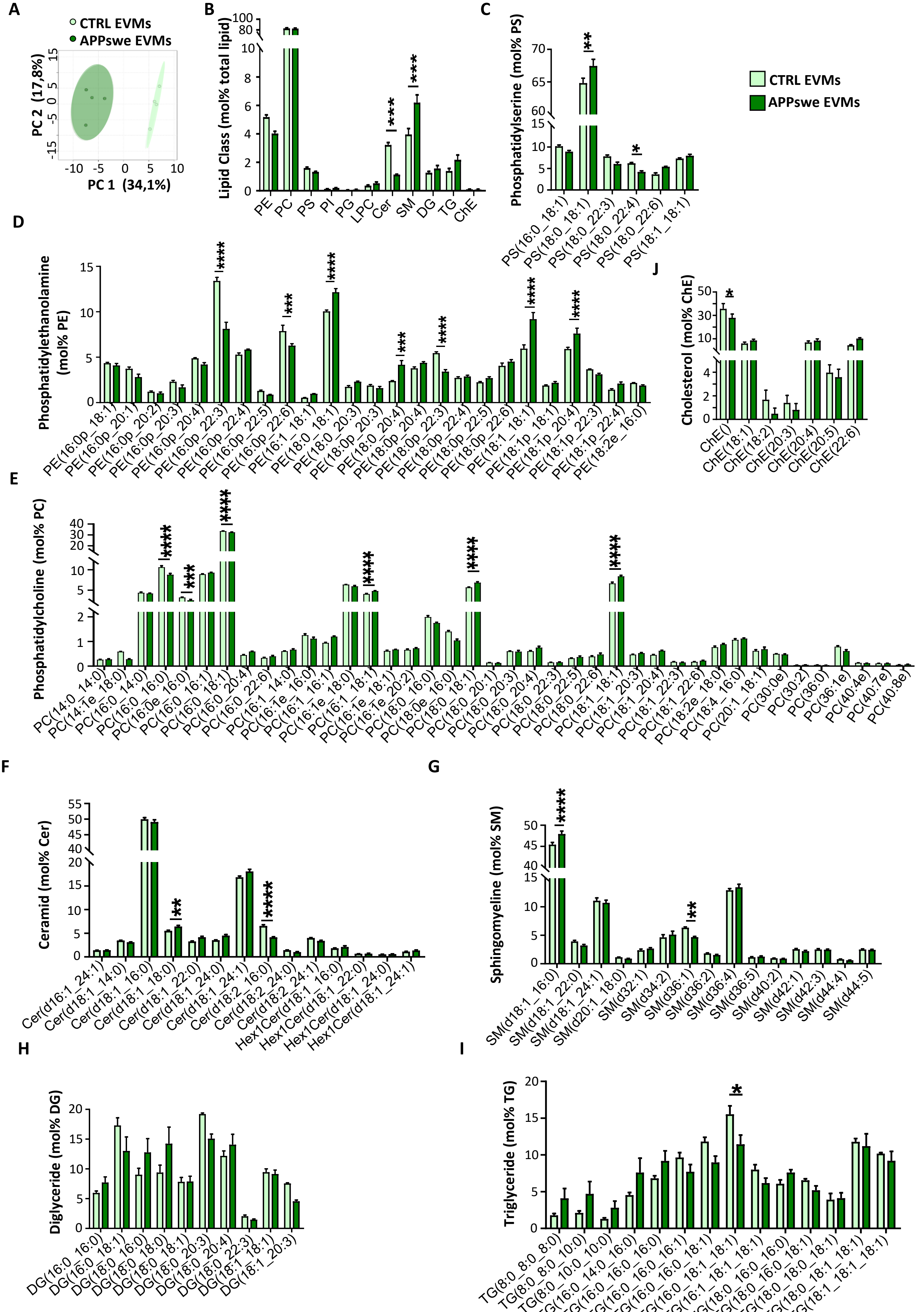


Eysert et al., Suppl. Figure 6

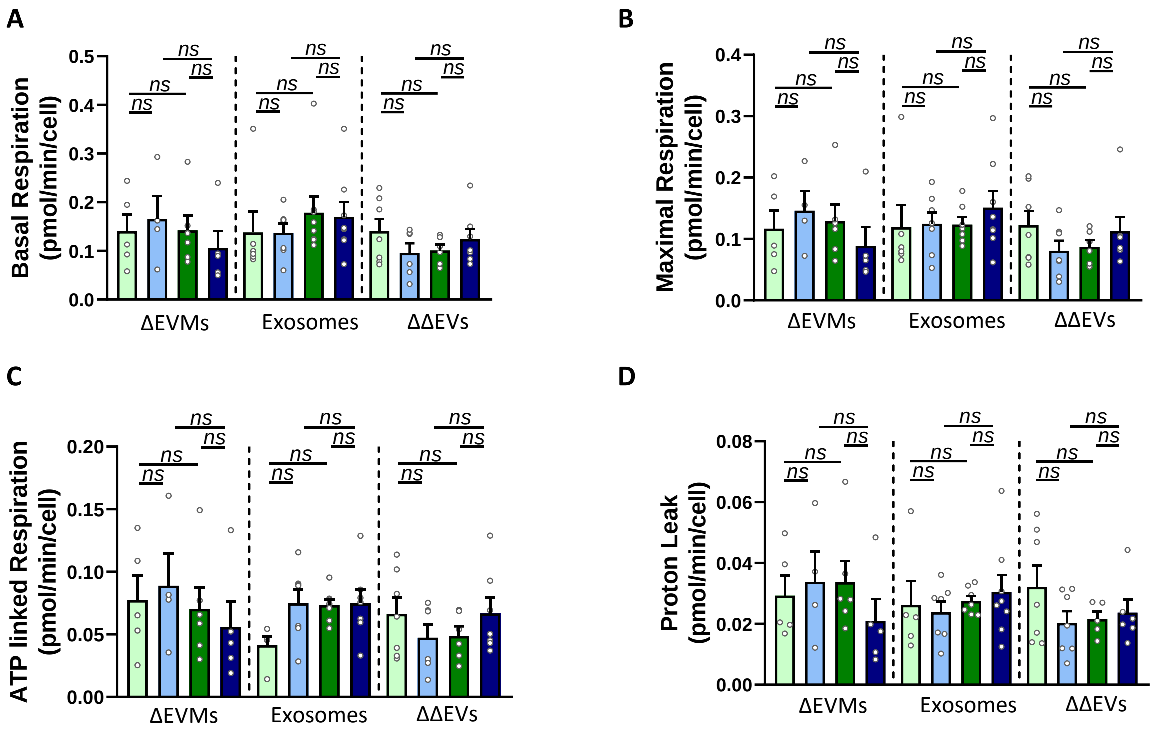


Eysert et al., Suppl. Figure 7
